# Effects of male age and female presence on male associations in a large, polygynous mammal in southern India

**DOI:** 10.1101/485144

**Authors:** P. Keerthipriya, S. Nandini, T.N.C. Vidya

## Abstract

We present a detailed study of male associations in a roving species, the Asian elephant, using six years of data on identified, nonmusth males. Adult males spent greater proportions of their time solitarily than in mixed-sex or in all-male groups. Old (over 30 years) males were sighted more frequently with their age-peers and less frequently with young (15-30 years) males than expected at random in all-male groups. Young males were not sighted more frequently with old males than with young males, and did not disproportionately initiate associations with old males. These results suggest that male associations, in the absence of females, primarily allow for old nonmusth males to test strengths against age-peers. Social learning from older individuals did not seem to be important in male associations, unlike that observed in African savannah elephants. We also found a constraint on the sizes of all-male groups, similar to that seen in female groups in our study population, and with male group sizes being smaller than that of African savannah elephants. However, most males had a significant top associate in female absence. In mixed-sex groups, male associations occurred at random, suggesting that males were tracking female groups independently. Thus, we find some differences in male social organisation compared to the phylogenetically related African savannah elephant that occupies a similar niche, and suggest that ecological factors might have shaped the differences in these male societies.

## Introduction

Adult males and females of many large mammals exhibit sexual dimorphism and strikingly different lifestyles, with female philopatry and male dispersal (see Greenwood 1980, Ruckstuhl and Neuhaus 2000). Following dispersal from their natal groups, males may spend some time alone or in all-male groups, and then form a) relatively stable bonds with female groups (for example, Hrdy 1977 in Hanuman langurs, Packer and Pusey 1987 in lions, Alberts and Altmann 1995 in baboons), b) form mixed-species groups during the breeding season and all-male groups outside of the breeding season (for example, Clutton-Brock *et al*. 1987 in red deer, Villaret and Bon 1995 in Alpine ibex, Mooring *et al*. 2003 in desert bighorn sheep), or c) rove between different female groups and form only temporary associations with female groups throughout the year (for example, Best 1979 in sperm whales, Poole 1982 in African savannah elephants, Desai and Johnsingh 1995 in Asian elephants, Baird and Whitehead 2000 in killer whales). The interactions between males themselves in polygynous species with these different kinds of lifestyles are expected to be competitive rather than affiliative, with males competing with one another for access to receptive females (van Hooff and van Schaik 1994). Therefore, strong associations are not expected between males in species with female-philopatry and may occur primarily in the context of coalitions to defend or to contest access to females (for example, Saayman 1971 in baboons, Connor *et al*. 1992, Gerber *et al*. 2020 in bottlenose dolphins, van Hooff and van Schaik 1994 in non-human primates, Wagner *et al*. 2008 in hyaenas). However, male-male interactions in all-male groups may be less competitive or aggressive than those within mixed-sex groups (Pusey and Packer 1987 in primates, Robbins 1996 in mountain gorillas). All-male groups may provide an opportunity to test strength against and assess competitors in a more relaxed setting (Bon *et al*. 1993 in mouflon sheep, Chiyo *et al*. 2011 in African savannah elephants). Male associations in all-male groups may also be motivated by the opportunities available for social learning from older, more experienced males (Evans and Harris 2008, Chiyo *et al*. 2011, 2012 in African savannah elephants, Bercovitch and Berry 2014 in giraffes). Increased efficiency in obtaining food resources (river otters, Blundell *et al*. 2002) and improved defense against predators (sperm whales, Curé *et al*. 2013) are also possible benefits from associating with other males, despite possible intrasexual competition. There has been little study on male association patterns in mammals overall, especially on those species that rove between female groups and do not form stable multimale-multifemale groups or large multimale groups in the non-breeding season. We, therefore, wanted to examine the extent and nature of male associations in such a roving species that is large, polygynous, and faces potentially high male-male competition and potential group size restriction, but is also phylogenetically related to a species with complex male association patterns.

Asian elephants (*Elephas maximus*) are polygynous, with males and females exhibiting different morphologies and adult lifestyles. Female society in this species is organised into clans that show fission-fusion dynamics (de Silva *et al*. 2011, Nandini *et al*. 2017, 2018), while pubertal males disperse from their natal groups and only temporarily associate with other males and with female groups thereafter (McKay 1973, Desai and Johnsingh 1995). There are often no clearly defined bull areas. Males are not known to form coalitions to defend females. Males can breed throughout the year, but females are sexually receptive only for a few days (Eisenberg *et al*. 1971) every four to five years, making receptive females a rare resource, for which males are expected to compete intensely. Male-male dominance interactions have been observed, indicating contest competition (McKay 1973, Daniel *et al*. 1987, Chelliah and Sukumar 2013, Keerthipriya 2018).

Male Asian and African elephants are thought to be more reproductively active during musth (a rut-like state); however, nonmusth males also acquire some mating and reproductive success (Hollister-Smith *et al*. 2007, Rasmussen *et al*. 2007, Chelliah and Sukumar 2015, Kabini Elephant Project, unpublished data). Therefore, competition for females may also exist among nonmusth males, and male-male affiliative associations are expected to be weak. However, males in the African savannah elephant have been shown to have complex relationships, with males preferring to associate with age-peers (Chiyo *et al*. 2011, Goldenberg *et al*. 2014 – in the case of sexually inactive males) and related males (Chiyo *et al*. 2011), and there is some evidence for older males being preferred associates or being more central to male society than young males (Evans and Harris 2008, Chiyo *et al*. 2011, Murphy *et al*. 2020). Male associations were also shown to facilitate social learning: bulls who had an older crop raider as a top associate were more likely to raid themselves (Chiyo *et al*. 2012). Thus, temporary all-male groups seem to provide an opportunity to spar and test strengths, and also possibly for younger males to learn from knowledgeable, older males in African savannah elephants.

Although Asian and African elephants were previously assumed to have similar societies, we now know that there are some differences between the female Asian elephant and African savannah elephant societies (de Silva and Wittemyer 2012, Nandini *et al*. 2018), probably because of a constraint on female group sizes due to ecology in the Asian elephant (Nandini *et al*. 2017, 2018). Since males, being larger and continuing to grow in size as they age (Sukumar *et al*. 1988), are likely to require more food than females, such a constraint might also exist in male Asian elephants and lead to differences in male societies across species, despite their phylogenetic closeness. Moreover, male African savannah elephants were known to return to the same bull areas (areas frequented by males and not by many female groups) when sexually inactive (Poole 1982), providing an opportunity for repeated associations with specific individuals (although the selection of a bull area may also possibly be a decision to associate with other males in the area; see Lee *et al*. 2011). An absence of separate bull areas in Asian elephant populations might reduce the frequency of males meeting each other, and hinder the formation of strong male associations. We, therefore, wanted to examine associations among adult male Asian elephants to find out whether possible ecological differences correlated with a different male social structure than in the African savannah elephant, despite the phylogenetic similarity between species.

We analysed associations of nonmusth adult males in this paper because musth males spend only a small proportion of their time in all-male groups (Keerthipriya *et al*. 2020). We aimed to examine the prevalence, strength, and stability of associations among nonmusth Asian elephant males and the factors (male age, presence of female groups, restriction on group size) that might affect male associations. We hypothesized that male (adult, nonmusth, male Asian elephants in this paper, unless specified otherwise) associations might be based on opportunities available for a) social learning from older individuals and/or b) testing strengths. Increased efficiency in obtaining food resources was not likely to be a factor leading to adult male groupings in elephants because individuals require large amounts of food and grouping is expected to create food competition. Defense against predators was also not likely to be important because adult male elephants do not have any natural predators and our work was carried out in Protected Areas. We did not examine genetic relatedness as a cause for associations.

We set out to address the following specific questions:

1. *What are the proportions of time that nonmusth males spend in all-male groups and mixed-sex groups and how are they related to male age?* While examining male associations regardless of the males’ musth status, we had found that age did not affect the time spent in mixed-sex or all-male groups (Keerthipriya *et al*. 2018). We had also found that old males increased their association with females when they entered musth, while young males decreased their association with females when they were in musth, compared to their nonmusth associations (Keerthipriya *et al*. 2020). Therefore, we expected that nonmusth older males would spend less time with female groups than nonmusth younger males. Although older males were seen more often than younger males in all-male groups of the African savannah elephant (see Chiyo *et al*. 2011, Goldenberg *et al*. 2014), that pattern might be reversed among nonmusth males in Kabini if there was a constraint on group size.
2. *How does male age and the presence or absence of females in the vicinity affect patterns of associations between nonmusth males?* We expected male age to affect associations differently depending on whether males associated primarily for social learning or for testing strength. We expected males to use their time in all-male groups, rather than in mixed-sex groups, to test strength. However, social learning from older males could occur in female absence (learning related to resources) and/or female presence (learning related to reproduction). If male associations were primarily based on social learning from older individuals, younger males would seek out older males more often than expected by chance alone. If male associations were primarily driven by opportunities for social learning, but older males were restricted in the amount of time they spent with other males (possibly due to group size constraints), older males might not spend more time in all-male groups, but the proportion of sightings of young males in which young males associated with older males would be higher than the proportion of sightings of young males in which young males associated with other young males. Older males would also be better connected in male networks if they were preferred as associates by young and old males, although better connectedness of old males could also be observed if old males had stronger age-peer associations than younger males. If the primary purpose of male associations was to test strengths, males would be expected to associate (relative to population age-structure) to a greater extent with age-peers than with much younger or older individuals, whose relative strengths are easily assessed by size differences. We did not have *a priori* expectations about whether old or young males would be more likely to associate amongst themselves if there were preferential associations with age-peers. It was possible that young males might show greater preferential association with age-peers than old males since old males might better know their strengths (through experience) than young males. However, it was also possible that old males might show greater preferential association with age-peers because, being more competitive breeders than young males, resolving dominance relationships might be more crucial for them. Since competition for females could play a major role in how males associated, we examined male associations in the immediate presence and absence of females. Unlike the case of the African savannah elephants, no clear indicators of active and inactive sexual states outside of musth have been recognised in Asian elephants. We expected the average number of other males that males met to be lower, and associations to be weaker, in the presence of females than in their absence. It should be noted that if males spent a greater amount of time in the absence of females than in their presence, males would likely also meet other males more frequently in female absence, resulting in the male association network in female absence being denser and better connected than that in female presence. However, this difference should not be significant after controlling for the difference in the number of sightings, if the greater number of associations was merely an effect of the greater amount of time spent. Due to the expected competition among males, we also predicted smaller experienced group sizes of the males in female presence than absence. It was also possible that males approached female groups by tracking them independent of other males, in which case, we expected males to associate with each other at random. In such a case, we had no *a priori* expectation about whether the group sizes of males would be higher in female presence or in female absence.
3. *Is there a restriction on male group size?* If there was a constraint on the group sizes of all-male groups, we expected a trade-off between the number of associates of a male and how often the focal male was sighted with those associates. We also expected older males to spend less time in all-male groups or to form smaller all-male groups than younger males because of greater food competition (due to larger body size). However, a similar pattern could also be observed if young males had stronger age-peer associations than old males. On the other hand, if there were stronger age-peer associations among old than young males, we would expect old males to spend more time in all-male groups or to form larger all-male groups than young males, although group-size restriction might reduce the difference.
4. *Are there preferential associations between nonmusth males and, if so, are they stable over time?* We did not have any *a priori* expectation about whether preferred, stable associations should be present or not, but, if they occurred, we expected them to be less frequent than that in the African savannah elephant due to possible group size constraints.

## Methods

### Field data collection

The field study was carried out in Nagarahole and Bandipur National Parks and Tiger Reserves (Nagarahole: 11.85304°-12.26089° N, 76.00075°-76.27996° E, 644 km^2^; Bandipur: 11.59234°-11.94884° N, 76.20850°-76.86904° E, 872 km^2^) in southern India from March 2009 to July 2014. Nagarahole and Bandipur National Parks are separated by the Kabini reservoir, and we refer to the elephants in the two parks as the Kabini population. Because of the high density of elephants around the reservoir and better visibility for behavioural observations, our sampling was centred around the reservoir, and extended to the forests in either direction with lower frequency of sampling (see Nandini *et al*. 2017). We sampled pre-selected forest routes (see Nandini *et al*. 2017 for details) in the study area from early morning to late evening (~6:30 AM am to 6:00-6:45 PM depending on field permits and light conditions). There were no distinct bull areas in our study area. The adult sex ratio in the study area was 1 male: 4-5 females (Gupta *et al*. 2016).

We tried to sex, age, and identify all the elephants sighted. Asian elephants are sexually dimorphic, with males being taller and bulkier than females apart from differences in genitalia. Females do not possess tusks, although some males (called makhnas) are also tuskless. We estimated age based on shoulder height, body length, skull size, and skin folds (see Vidya *et al*. 2014), with semi-captive elephants in the same area serving as a reference for ageing older animals. We placed males into the following age categories, based on their estimated ages: calves (<1 year), juveniles (1-<5 years), sub-adults (5-<15 years), young adults (15-<30 years), and old adults (>=30 years). Young adult males were likely to have completely dispersed from their natal herds but were possibly less reproductively competitive than old adult males, based on studies of African savannah elephants (Poole 1982, Poole *et al*. 2011). The ages used for classifying males into these categories were those calculated at the mid-point of the study period (November 2012). We identified individuals based on a combination of ear, back, tail, tusk, and body characteristics (detailed in Vidya *et al*. 2014). We recorded group size, GPS location, time of sighting, and whether adult males were in the presence or absence of females. Females were classified as adults when they were 10 years old (see Nandini *et al*. 2018). Adult males were said to associate with a female group (one or more adult females and their young that were in close proximity and showed coordinated movement; see Nandini *et al*. 2018) if they fed within 10 m (easy physical reach) of a group member or interacted with any group member. When two males associated with the same female group at the same time, they were said to be associating with each other in female presence. Rarely (only three different sightings), males were seen to associate with subadult females (5-10 years old) in the absence of an adult female and this was also considered to be association in female presence. Males were said to associate with each other in female absence if they fed within about 50 m of each other and there were no females in the vicinity. At this distance, the males would be able to display or react to visual signals, apart from sensing one another through sound or smell. Males could indulge in sparring during their associations, but if males, upon encountering each other, displayed only aggressive interactions and moved away, they were not considered to be associating.

### Data Analysis

Data analysis was carried out using only those sightings in which all adult males were aged and identified and female group compositions (if applicable) were known. Of the 878 days of field work between 2009 and 2014, elephants were sighted on 853 days, identified adult males were sighted on 718 days, and nonmusth adult males on 681 days. In many of the analyses mentioned below, only males who were sighted at least 10 times in that particular category (such as group composition type or female presence) while not in musth were used, as associations of males seen rarely are unlikely to represent their actual association patterns and may bias the results. Similarly, if there was a comparison between different categories (such as associations in female presence and absence), common males sighted at least 10 times (when not in musth) in each of the categories were used, unless otherwise mentioned. ANOVAs, GLMs/GLMEs (General Linear Models/Generalised Linear Mixed-Effects Models), and non-parametric tests were performed using Statistica 7 (StatSoft, Inc. 2004) and randomisations (as randomisations of observational data can account for potential sampling bias in social network analyses; see Farine 2017) were carried out using MATLAB (MATLAB R2011a, MathWorks, Inc, 1984-2011, www.mathworks.com) unless specified otherwise.

### Proportions of time spent in all-male and mixed-sex groups and their relationship with male age

We calculated the number of minutes individual males (that were seen on at least 5 different days when not in musth) were seen in the following group types and calculated the proportions of each individual’s time spent in such groups: 1) solitary, 2) all-male groups with only one adult male (but including subadult or juvenile males and, therefore, not solitary), 3) all-male groups with more than one adult male, and 4) mixed-sex groups. Since the four proportions add up to one and are, therefore, not independent, and the number of males seen in group type 2 was small, we compared only two of the four categories – all-male groups with more than one adult male and mixed-sex groups. Males were categorized into old and young adults as explained in the *Field data collection* section above. We carried out Wilcoxon’s matched-pairs tests, separately on old and young males, to test whether the proportions of time spent in the different group types differed significantly. Since the proportions were calculated for each male, each male was represented only once in this analysis, effectively making male identity random. In order to find out whether old males spent a smaller proportion of their time in mixed-sex groups than young males did, we used a Mann-Whitney *U* test.

### Changes in male associations and time to sighting independence

Using data from the last two years of observation (2013-2014), we examined continuous observations of individual males and recorded the number of changes in that male’s associations in five-minute bins (all observations that lasted less than five minutes were removed for this calculation). If there were changes in the identities of the adult males in the focal male’s group or if the focal male’s association changed from female presence to absence (or vice versa), they were counted as changes in the focal male’s associations. The time interval at which there was roughly equal probability of a male’s association changing or not would indicate the time interval at which sightings could be recorded as independent of each other (in order to avoid pseudoreplication of data). We found this time interval to be about 80 minutes (see Supplementary Material 1). Therefore, for all subsequent analyses of sighting data, we only included independent sightings based on this time interval.

### Effect of male age and the presence or absence of females on male association patterns

#### Male age and social learning

We examined the initiation of associations and the pattern of associations between males to find out whether social learning might be a reason for associating. As mentioned in the Introduction, if associations were primarily based on social learning, younger males would be expected to seek out older males more often than expected by random chance. We, therefore, examined all the instances in which two males (dyadic combinations in the small number of cases where there were more than two males) of different ages were in close proximity and one approached the other and associated with it. It was possible under such circumstances for either of the males to approach the other. We randomly chose one of the males as the focal male and used a generalized linear mixed-effects model (GLME) with a binomial dependent variable (0 if the focal male approached the associate and 1 if the focal male was approached by the associate), age difference (age of the focal male minus the age of the associate) as a continuous predictor variable, and dyad identity as a random factor. If younger males approached older males more often than expected, a significant positive effect of the age difference on the dependent variable would be found. We used the *fitglme* function in MATLAB R2011a, with a logit link function and Laplace estimation method, and the model was fit using maximum likelihood. For this analysis, we used data only from the years 2011-2014, during which detailed behavioural observations were available, and did not include dyads that were already present when we began the observation. We only used instances (of all males) when observers were present during the beginning of the association and one of the males clearly moved towards the other.

We had expected that if social learning was the primary reason for the formation of all-male groups, but older males were restricted in the amount of time they spent with other males, the proportion of young males’ time (sightings) that was spent with older males would still be higher than the proportion of young males’ sightings spent with other young males. Therefore, we compared these proportions, in female presence and absence, using Wilcoxon’s matched-pairs tests. (Young males would spend a greater proportion of their sightings with other young males than with old males if the testing-strength hypothesis was correct.)

We had also expected older males to be better connected in male networks if they were preferred as associates by both young and old males (or disproportionately by old males). We examined this by analysing social networks of males. We first calculated the association index (AI; Ginsberg and Young 1992) between pairs of identified males as the number of times (independent sightings) two males were sighted together (*N*_AB_) divided by the total number of (independent) sightings of the two males. AI between each pair of males was calculated separately in female presence and female absence (for instance, AI_AB(F_abs)_=*N*_AB(F_abs)_/(*N*_A(F_abs)_+*N*_B(F_abs)_-*N*_AB(F_abs)_, where *N*_A(F_abs)_ is the total number of sightings of A in female absence and *N*B(F_abs) is the total number of sightings of B in female absence). We used these AIs to calculate association strength (or weighted degree) for each individual male, as the sum of the AI values between a male and all his associates. We calculated the difference in association strength between the two age-classes (mean association strength of old males – mean association strength of young males) in the observed dataset. We then obtained the same value from each of 5000 permuted datasets (see the section below for details of permutations) and compared the difference in the observed dataset with those from the permuted datasets (see section below; *P*<0.025 for statistical significance as we had no prior expectation about whether the observed values would be lower or higher than the permuted values).

#### Male age and testing strengths

In order to find out whether males preferentially associated with age-peers (as predicted by the testing-strengths hypothesis) more often than expected by chance, we used a permutation procedure following Whitehead (2008, pg. 124). This procedure was carried out on the datasets of males in female absence. We generated 5000 *males permuted* datasets, by flipping (switching) adult males across independent sightings, while keeping the group size and the number of sightings for each male constant, in each permuted dataset. The number of flips performed in each permutation was five times the number of sightings in that dataset. We calculated the number of times old (>=30 years) and young (15-30 years) males were sighted with other males of the same or different age class in the observed dataset and compared these observed values with the values from the permuted datasets. We used the distribution of values from the permuted datasets to determine whether the observed value was greater or lower than that expected by chance (*P* = proportion of permuted values that were higher – if the observed value was higher than more than half of the permuted values – or lower – if the observed value was lower than more than half of the permuted values – than the observed value; *P*<0.025 for statistical significance).

We also compared the proportion of their sightings that old males spent associating with other old males with the proportion of their sightings that they spent associating with young males using Wilcoxon’s matched-pairs tests. A higher proportion of sightings with other old males than young males would be seen if the testing-strengths hypothesis was correct.

### Male associations in the presence and absence of females

We performed a similar permutation procedure as described in the section above on the female presence dataset, comparing the observed numbers of sightings in which males associated with their age-peers and the other age-class. If association between males was influenced by competition for females, we expected old males to associate less frequently than that expected by random chance. We had also expected the male association network to be denser and better connected in female absence than in female presence because of potential competition between males in the presence of females. We, therefore, compared the following network statistics between male association networks in female presence and absence: mean *degree* (number of associates of a focal male), mean *clustering coefficient* (the proportion of the total possible connections between a male’s associates that exist), mean *path length* (the smallest number of connections between two individuals in the network), and *network density* (the proportion of all possible connections that exist in the network) (Latapy 2008; see Wasserman and Faust 1994). We compared these network statistics and mean AIs between female presence and absence using a sampled randomization test (Sokal and Rohlf 1981, pp. 791-794). We created 5000 permuted datasets (permuted by randomly assigning sightings to female presence or absence, while conserving the sample sizes for both the categories and ensuring that at least ten sightings of each male were assigned to each category in each permutation). We then compared the observed differences in network statistics and AI between the original female presence and female absence datasets with the differences in statistics between the permuted ‘female presence’ and ‘female absence’ datasets. The probability of a significant difference between the observed values was calculated as the proportion of permutations in which the difference in statistic between the permuted datasets was greater than or equal to the difference in statistic between the observed datasets. Since the total numbers of sightings in female presence and female absence were conserved, a significant difference in this test would indicate a difference in network statistic that could not be attributed to the difference in the numbers of sightings.

We compared the degree distributions of association networks in female presence and absence with their Poisson expectations (expected for a Erdös-Rényi random network; Erdös and Rényi 1960) to test whether males were associating with each other at random.

We examined the effect of female presence on male group size by performing a GLM on experienced group sizes (experienced by individual adult males; counted as the number of adult males in each sighting of the focal male, including the focal male). We used female presence/absence as a fixed factor and male identity as a random factor.

### Restriction on all-male group size

If there was a constraint on group size in all-male groups, a negative relationship would be seen between the number of associates of a male and how frequently the focal male was sighted with those associates. We, therefore, calculated the average of the proportions of sightings of a focal male (out of all the sightings of that male, including solitary sightings) that the male spent with his different associates. This average proportion of the focal male’s sightings spent with his associates was correlated with the number of associates of that male (in female absence) using a Spearman’s rank-order correlation. We also examined whether old males spent less time than young males in all-male groups by using the data on proportions of time spent in different types of groups (section towards the beginning of the Methods) and performing a Mann-Whitney *U* test. We also examined the relationship between the mean experienced group size of a male and his age using Spearman’s rank-order correlation.

### Preferred male associations and stability of associations

We tested for preferred associations or avoidance amongst identified males across sampling periods smaller than the entire dataset using SOCPROG 2.6 (Whitehead 2009). We used a sampling period of 14 days and 10,000 permutations with 10,000 flips for each permutation. We used the ‘*permute associations within samples*’ method, which tests for long-term (across sampling period) preferences and avoidances (Whitehead 2009). The presence of long-term preference/avoidance is indicated by significantly (>95% of the values of the randomised datasets) higher SD and CV of AI values from the real dataset when compared to the randomised datasets. We used sightings of all males for this analysis.

We additionally determined a top associate (based on AI value) for males, separately in female presence and in female absence. Directed networks of males and their top associates were constructed using Gephi 0.8.2 (Bastian *et al*. 2009). In order to examine whether the identity and strength (AI) of a male’s association with his top associate was different from random, we again used the procedure for permuting associations (Whitehead 2008, pg. 124) explained in a section above. We permuted associations between males separately in the female presence and female absence datasets (5000 permutations on each dataset, with the number of flips in each permutation being five times the number of sightings in that dataset). We compared the AI of a male and his observed top associate with AI values for the same dyad from the permuted dataset, and considered the association to be significant if the observed AI was greater than 95% of the permuted values. In each permutation, for each male, we also recorded the identity of his permuted top associate. We examined how often the male had the same top associate in the permuted datasets, and considered the identity of his top associate to be significantly different from random if the observed top associate was not the top associate in greater than 95% of the permuted datasets.

In order to find out whether male associations were stable across years, we compared AI matrices between consecutive years, using those males that were common to and sighted at least five times in both years, by performing Mantel tests of matrix correlation (Mantel 1967), with 5000 permutations, in MATLAB (MATLAB R2011a, MathWorks, Inc, 1984-2011, www.mathworks.com). Due to limitations of the number of sightings, only data from 2011-2014 and only in female absence were used for this analysis.

## Results

### Proportions of time spent in all-male and mixed-sex groups and their relationship with male age

We sighted 83 nonmusth males in all (see Supplementary Material 2), of which 44 were seen in the presence of females and 81 in the absence of females. Based on the set of nonmusth males seen on at least five different days (*N*=38), we found that males spent an average (± SD) of 12.3% (± 11.79%) of their time in all-male groups and 29.6% (± 22.73%) of their time in mixed-sex groups (Figure 1a). Old males spent similar proportions of their time in all-male and mixed-sex groups (Wilcoxon’s matched-pairs test: *T*=63.00, *Z*=0.259, *N*=16, *P*=0.796), whereas young males spent a significantly higher proportion of their time in mixed-sex groups than in all-male groups (Wilcoxon’s matched-pairs test: *T*=9.00, *Z*=3.815, *N*=22, *P*<0.001, see Figure 1b). Old males spent a significantly smaller proportion of their time in mixed-sex groups than young males did (Mann-Whitney *U* test: *U*=79.50, *Z*adj=-2.853, *P*=0.004, see Figure 1b).

**Figure 1.**
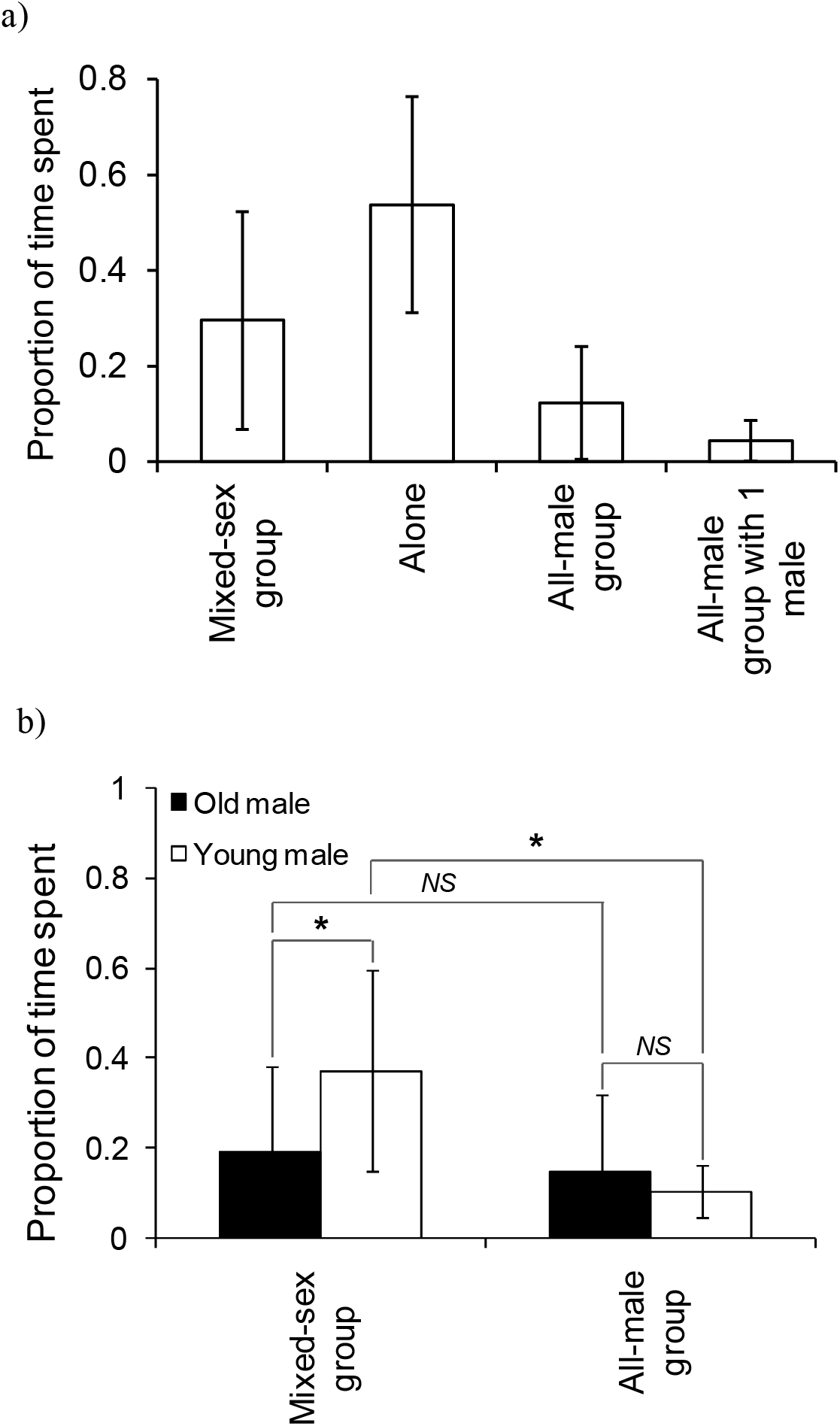
The proportion of time that a) males (sighted on five days or more) spent in different group types and b) old and young males (sighted on five days or more) spent in mixed-sex and adult all-male groups. Pairwise comparisons with significant results are marked with an asterix, and those with non-significant results are marked as *NS* (Not Significant). Error bars are standard deviations.

### Effect of male age and the presence or absence of females on male association patterns

Based on the 80-minute cutoff, there were 2466 independent sightings of adult nonmusth males (374 sightings in female presence and 2092 sightings in female absence) in the study period. Fourteen males (4 old and 10 young males) were sighted at least 10 times in female presence, and 32 males (13 old males and 19 young males) were sighted at least 10 times in female absence. Only 12 males were sighted at least 10 times each in the presence and absence of females.

#### Male age and social learning

The age difference between the focal male and the other male did not significantly affect who approached whom either in female presence (GLME: *Estimated coefficient_Age difference_*= - 0.035, *t*_stat_= −0.821, *N*=19 approaches, *df*=17, *P*=0.423) or in female absence (GLME: *Estimated coefficient_Age difference_*= −0.026, *t*_stat_= −0.772, *N*=48 approaches, *df*=46, *P*=0.444) (Figure 2). Thus, old and young males were equally likely to approach each other to initiate association. We also found no significant difference in the proportion of its sightings that a young male spent with old males (Female presence: mean ± SD 0.058 ± 0.035; Female absence: 0.073 ± 0.086) and the proportion of its sightings a young male spent with other young males in female presence or absence (Female presence: mean ± SD: 0.111 ± 0.090; Wilcoxon’s matched-pairs test: *T*=6.50, *Z*=1.610, *N*=10, *P*=0.107; Female absence: 0.065 ± 0.045; Wilcoxon’s matched-pairs test: *T*=58.00, *Z*=0.114, *N*=19, *P*=0.910).

**Figure 2.**
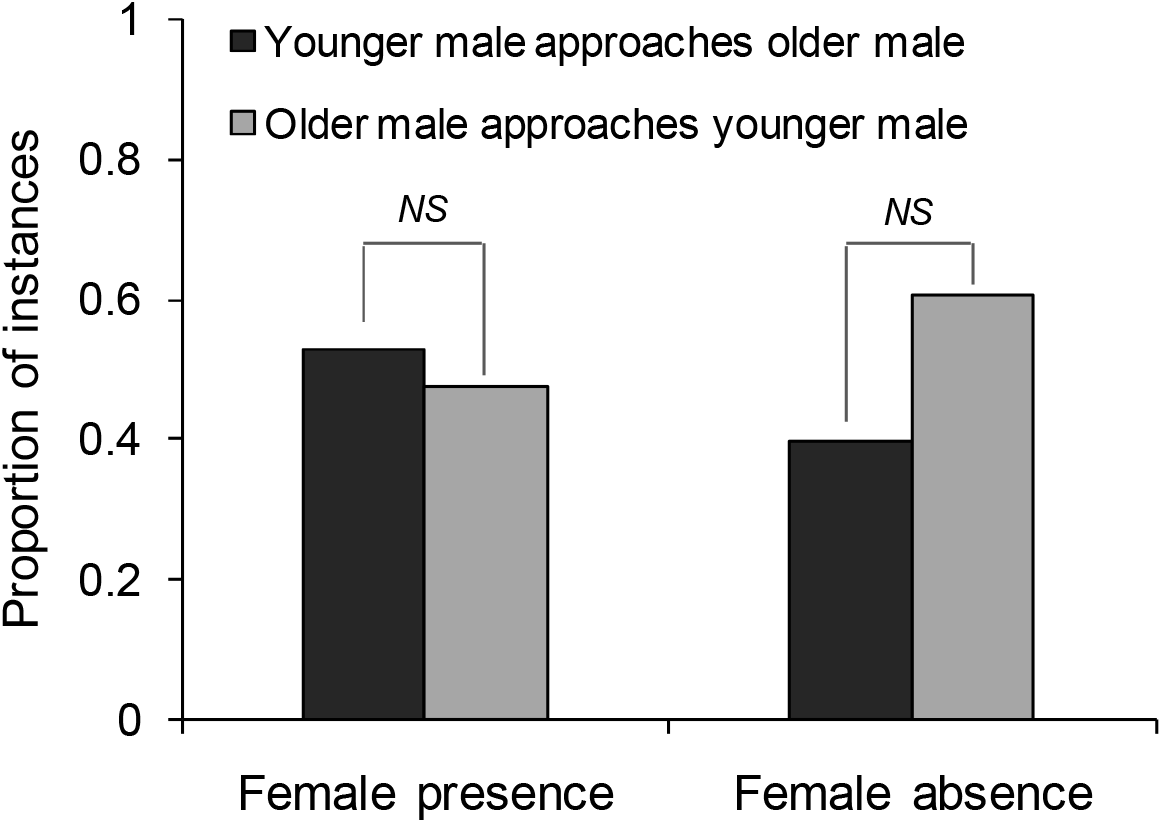
The proportion of instances when the younger male approached the older male, and the proportion when the older male approached the younger male, to initiate association, in female presence and absence. The proportions were not different from one another in female presence and absence and hence, pairwise comparisons results are marked as *NS* (Not Significant).

Examining the effect of male age-class on association strength, we found old males to have only a slightly higher association strength (mean=0.087) than young males (mean=0.052) in the absence of females (difference in mean association strength (old-young): observed dataset: 0.035; permuted datasets: mean (2.5 and 97.5 percentiles): 0.003 (−0.015, 0.020), *P*<0.001, Table 1). The observed difference in mean association strengths between the age-classes was not different from the permuted values in female presence (see Table 1).

**Table 1.**
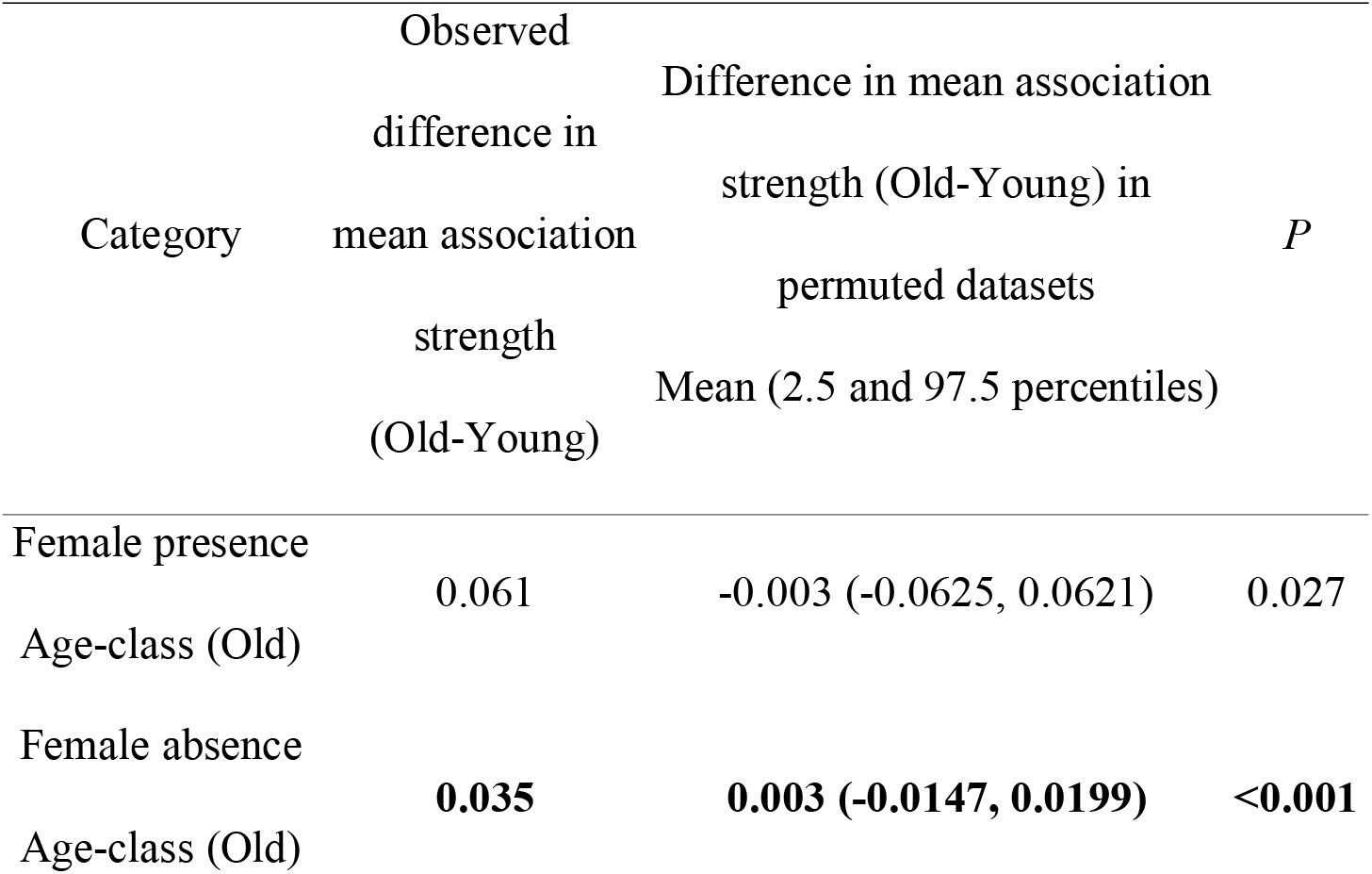
Difference in mean association strength between males of the two age-classes in observed and permuted datasets. The associated probability values (*P*=number of permutations whose difference in mean association strength is higher than that based on the observed dataset / number of permutations) are provided. Significant comparisons (*P*<0.025) are marked in bold.

| Category | Observed<br>difference in<br>mean association<br>strength<br>(Old-Young) | Difference in mean association<br>strength (Old-Young) in<br>permuted datasets<br>Mean (2.5 and 97.5 percentiles) | $P$ |
| --- | --- | --- | --- |
| Female presence<br>Age-class (Old) | 0.061 | -0.003 (-0.0625, 0.0621) | 0.027 |
| Female absence<br>Age-class (Old) | **0.035** | **0.003 (-0.0147, 0.0199)** | **<0.001** |

#### Male age and testing strengths

In female absence, old males were sighted together more frequently than expected by chance (permutation test: observed: 42 sightings, *males permuted* mean (2.5, 97.5 percentiles): 25.5 (18, 34) sightings, *P*<0.001; Figure 3a). Old and young males were sighted together less frequently than expected by chance (observed: 51 sightings, permuted: 69.4 (58, 80) sightings, *P*=0.001) and young males were sighted together as expected by chance (observed: 38 sightings, permuted: 42.9 (33, 53) sightings, *P*=0.178; Figure 3a). Although these tests were based on small numbers of males, the results remained unchanged when we repeated the analysis using all the identified males (Supplementary Material 3).

**Figure 3.**
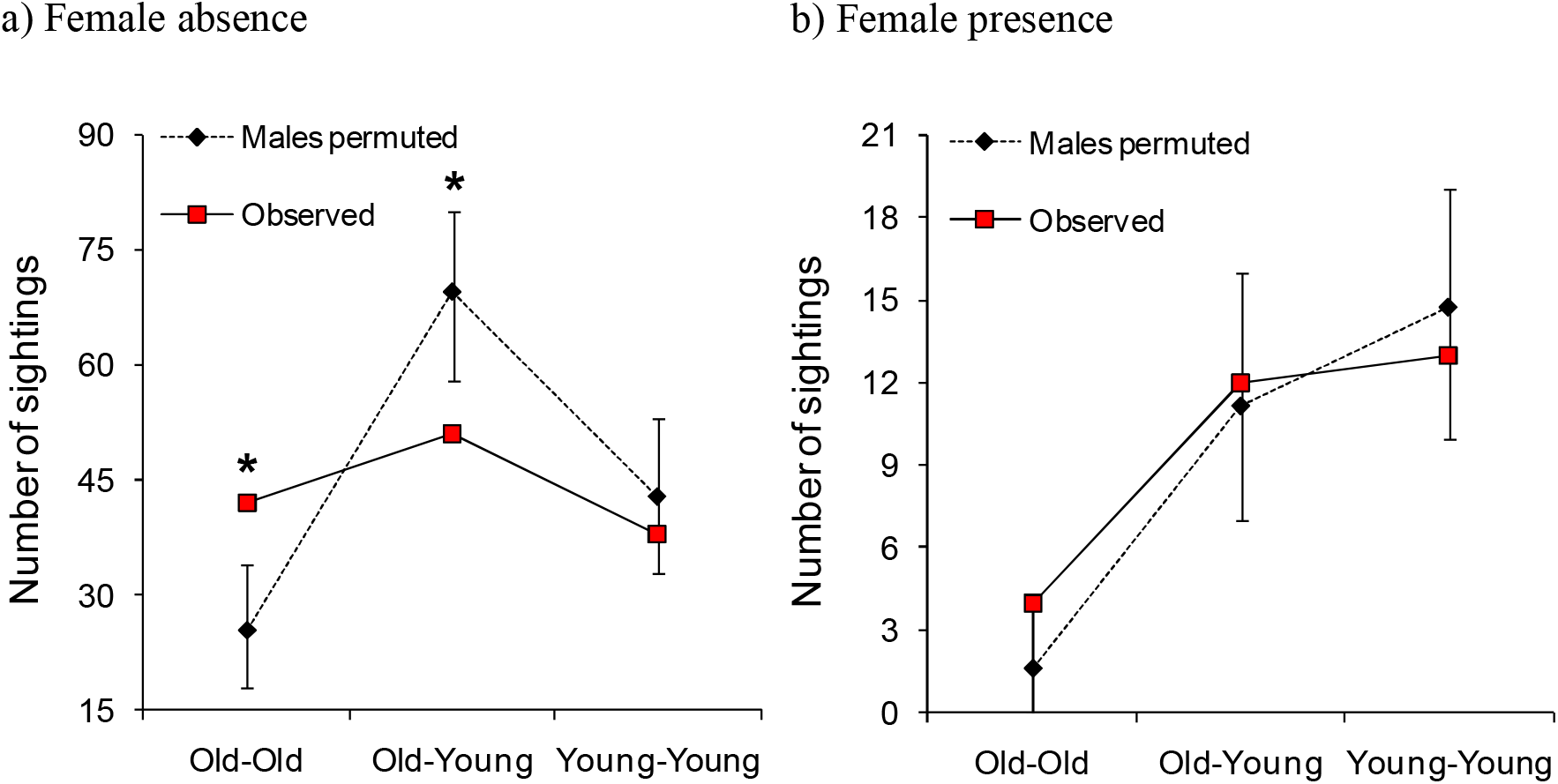
Permuted and observed numbers of times adult males (sighted 10 times or more in that category) of the same and different age-classes were sighted together in a) female absence and b) female presence. Comparisons where the observed value was significantly different from those of the permuted datasets are marked with an asterix. Old males are >=30 years and young males are 15-30 years old. Please note that the Y axis is on different scales in the two panels.

The proportion of their sightings that old males spent with other old males (mean ± SD: 0.141 ± 0.116) was higher than the proportion that they spent with young males (mean ± SD: 0.085 ± 0.069; Wilcoxon’s matched-pairs test: *T*=5.00, *Z*=2.489, *N*=13, *P*=0.013), in female absence.

### Male associations in the presence and absence of females

We found that old males were sighted together as expected by chance in the presence of females (permutation test: observed: 4 sightings, *males permuted* mean (2.5 and 97.5 percentiles): 1.6 (0, 4) sightings, *P*=0.069). The numbers of sightings were not different from those expected by chance in the case of old and young males seen together (observed: 12 sightings, permuted: 11.1 (7, 16) sightings, *P*=0.439) and young males seen together either (observed: 13 sightings, permuted: 14.7 (10, 19) sightings, *P*=0.289) (Figure 3b). These results remained unchanged when we repeated the analyses using all identified males without a sighting cutoff (Supplementary Material 3). All three age-class combinations of males were sighted together more often in female absence than in female presence (Figure 3). However, while the observed male association network appeared more connected and denser (higher mean degree, mean clustering coefficient, and density, and lower mean path length) in female absence than in female presence, none of these statistics was significantly different based on sampled randomisation tests (all *P*>0.05, 12 males seen at least 10 times each in female presence (249 sightings) and in female absence (855 sightings), Table 2). There was also no significant difference in mean AI between males in the presence and absence of females (Table 2, see Supplementary Material 4 for distributions). Thus, the seemingly more connected network in female absence was probably a result of greater time spent by males in female absence.

**Table 2.** Network statistics based on observed and permuted male associations in female presence and female absence. *P*=(number of times difference_random_ ≥ difference_observed_) / number of randomisations (5000). None of the comparisons was significant.

| Category | Mean AI | Mean degree | Mean clustering coefficient | Mean path length | Density |
| --- | --- | --- | --- | --- | --- |
| Female presence observed | 0.005 | 2.167 | 0.300 | 2.345 | 0.197 |
| Female absence observed | 0.003 | 4.500 | 0.577 | 1.697 | 0.409 |
| Female presence permuted mean (SD) | 0.004 (0.0011) | 1.463 (0.4246) | 0.224 (0.2263) | 2.054 (0.4013) | 0.133 (0.0386) |
| Female absence permuted mean (SD) | 0.003 (0.0002) | 4.887 (0.3184) | 0.592 (0.0723) | 1.623 (0.0621) | 0.444 (0.0289) |
| *P* value | 0.1126 | 0.9948 | 0.6316 | 0.2784 | 0.9448 |

The association network of adult males in female presence was not significantly different from a random network (*χ^2^*=6.122, *df*=8, *P*=0.633), but the network in female absence was significantly different from random (*χ^2^*=184.647, *df*=15, *P*<0.001) (see Supplementary Material 5).

Group sizes (number of nonmusth adult males in the sighting) were small in general, with the modal group size of all groups that had an adult male being 1 and the modal group size of all multi-male groups being 2 (Figure 4a). The experienced group sizes of common males (*N*=12, seen 273 times in all in the presence of females and 1304 times in all the absence of females) were larger in female presence (mean ± SD: 1.6 ± 0.80) than in female absence (mean ± SD: 1.2 ± 0.52; *F*_1,1553_=48.520, *P*<0.001). There was a significant main effect of male identity (*F*_11,1553_=2.918, *P*=0.045) and a significant interaction effect between male identity and female presence (*F*_11,1553_=2.157, *P*=0.014; Figure 4b).

**Figure 4.**
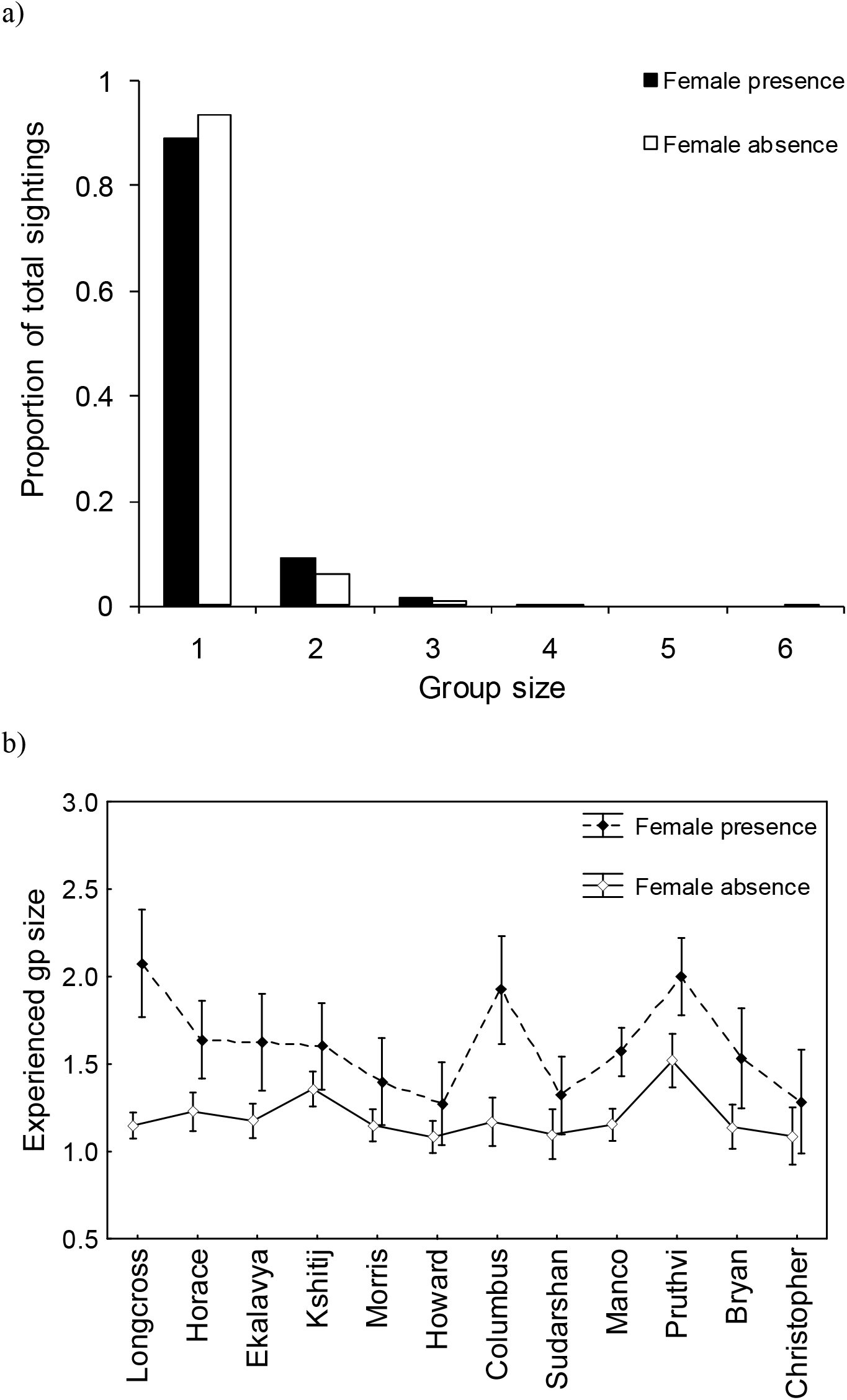
a) Proportion of sightings of all groups containing at least one male, of different group sizes (different numbers of adult males) and b) the experienced group sizes of common males in female presence and absence. Error bars are 95% CI.

### Restriction on all-male group size

The mean proportions of sightings that a male spent with an associate was significantly negatively correlated with the number of associates of that male (Spearman’s rank-order correlation: *N*=31 males, each with at least one associate and seen at least ten times in female absence, *r_s_*= −0.582, *P*<0.001; Figure 5). The relationship was even more negative when the mean proportion of all-male group sightings (rather than all sightings) spent with an associate was correlated with the number of associates of those males (*r_s_*= −0.984, *P*<0.001; Figure 5). Thus, in female absence, males that associated with a greater number of males spent a smaller proportion of their sightings on average with each of their associates. However, contrary to expectation, old and young males did not spend significantly different proportions of their time in all-male groups (Mann-Whitney *U* test: *U*=157.50, *Z*_adj_=0.547, *N*_Old_=16, *N*_Young_=22, *P*=0.584), and there was no significant relationship between male age and mean experienced group size (Spearman’s rank-order correlation: *N*=31 males, each with at least one associate and seen at least ten times in female absence, *r_s_*=0.213, *P*>0.05).

**Figure 5.**
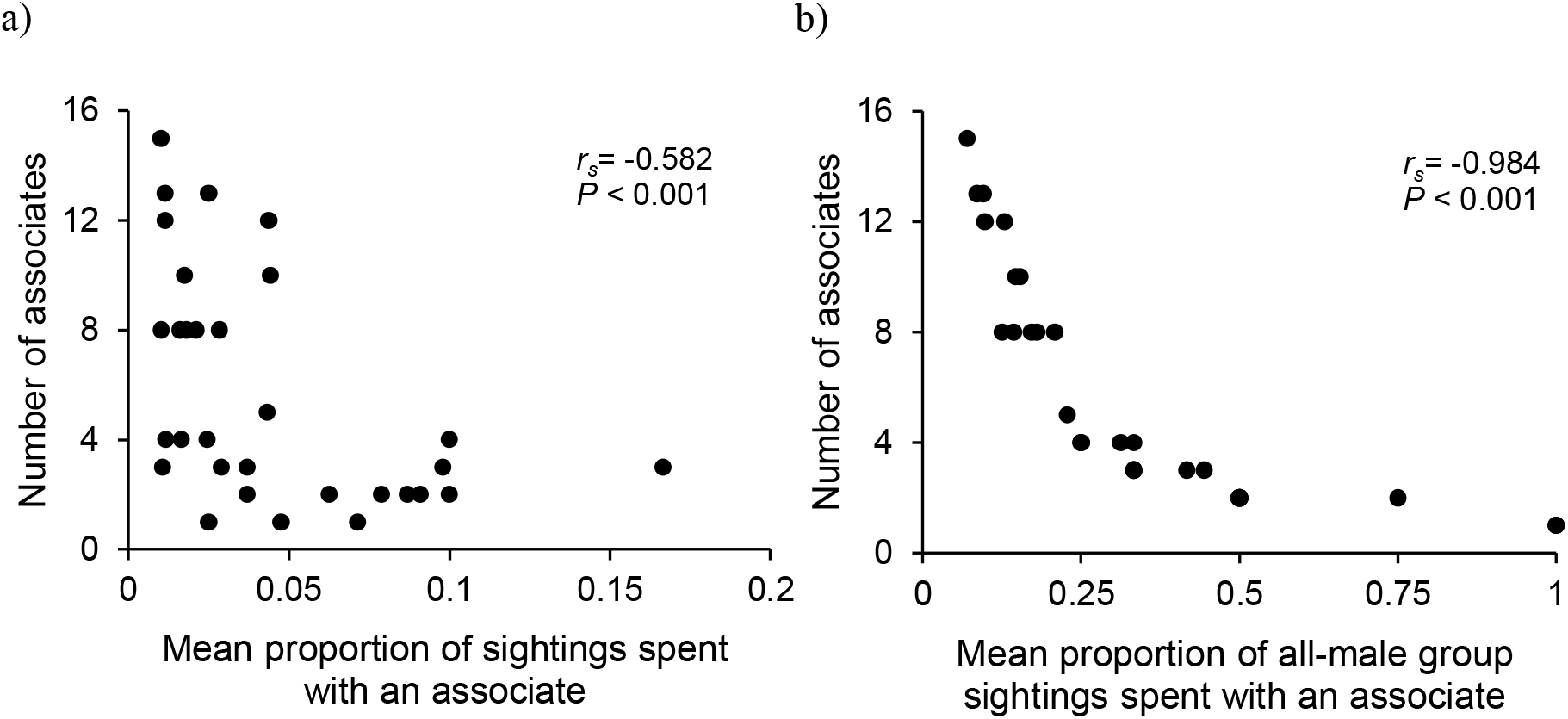
The mean proportion of a) sightings (out of the total sightings of the male) and b) all-male group sightings (out of the total all-male group sightings of the male), that a male spent with an associate plotted against the number of associates of that male. Results from Spearman’s rank-order correlations are written in the plot.

### Preferred male associations and stability of associations

We found no evidence of preferred male associations across 14-day sampling periods in female presence or in female absence (details in Supplementary Material 6). Only 14.3% (2 out of 14 males; 2 out of 15 dyads as one male had two top associates) of the males had a significant association with their top associates in female presence, whereas 61.3% (19 of 31 males; one male did not associate with any of the other males) of the males had a significant association with their top associate in female absence (mean significant AI with top associate in female absence=0.032, see Supplementary Material 4 for AI distribution; networks shown in Figure 6). We found that the identity of the focal male-top associate dyad was significantly different from random in 13.3% (2 out of 15) of the dyads in female presence and in 83.9% (26 out of 31) of the dyads in female absence.

**Figure 6.**
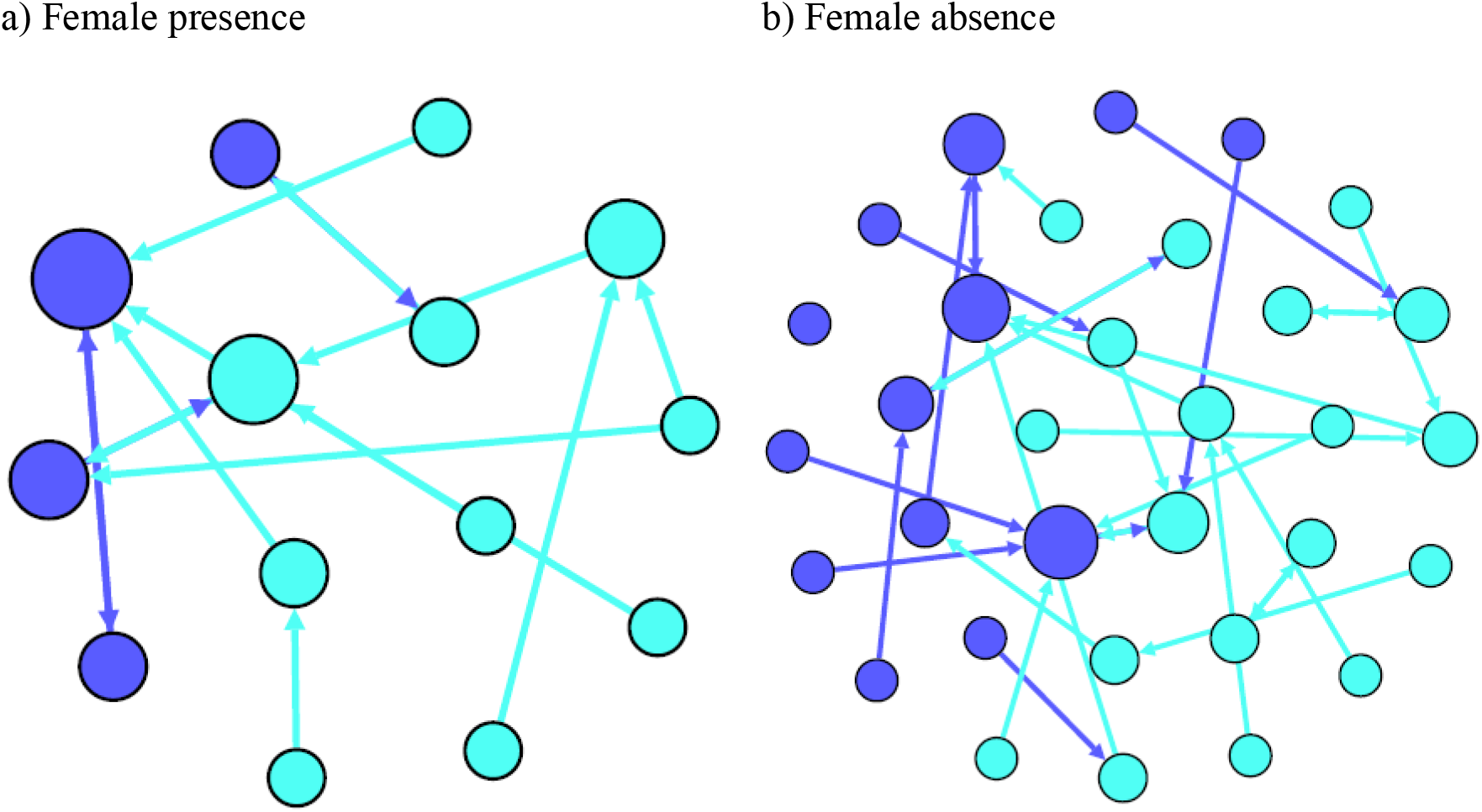
Networks of males and their top associates in a) female presence and b) female absence. Nodes representing old males are coloured dark blue and those representing young males are coloured light blue.

Mantel tests showed no significant correlation between association matrices across years, in two out of the three comparisons (Supplementary Material 7).

## Discussion

This is the first detailed study of nonmusth adult male associations in Asian elephants in a relatively undisturbed natural habitat. We found male-male affiliative associations to be weak as expected, and in contrast to that found in the African savannah elephant.

### Proportions of time spent in all-male and mixed-sex groups and their relationship with male age

Nonmusth adult males spent only ~12% of their time in all-male groups in Kabini, in contrast to ~63% after adjusting for age-effects (Chiyo *et al*. 2011) and 30.5% of all bull sightings (Lee *et al*. 2011) in the Amboseli African savannah elephant population. Unlike Kabini, Amboseli has distinct “bull areas” (Poole 1982) that ‘sexually inactive’ adult male elephants inhabit but females seldom do, and it is possible that adult males may be more likely to encounter and form strong associations with one another in such bull areas. However, Chiyo *et al*. (2011) used observations of all-male groups of males in an area used by both males and females; therefore, the difference cannot be attributed to bull areas in that case. We found evidence for a restriction of all-male group sizes in Kabini (see below), and this restriction may contribute to the smaller time spent in all-male groups. We expected and found old nonmusth males to spend less time with female groups than young nonmusth males. Whereas old males spent a similar proportion of their time in all-male and mixed-sex groups, young males spent more of their time in mixed-sex than all-male groups. As young males invest less in a competitive breeding strategy like musth (see Keerthipriya *et al*. 2020), their greater associations with females might facilitate learning about female groups in areas they have dispersed to and possible opportunistic matings (see Chelliah and Sukumar 2015, Kabini Elephant Project, unpublished data), or reflect a reduced cost of feeding alongside females compared to that faced by old males. Further data are required to examine these possibilities.

### Effect of male age on association patterns: Reasons for adult male associations

As adult Asian elephants do not face any major risks of predation within protected areas and are not known to form coalitions, we had hypothesised that all-male groups could provide an opportunity for younger males to learn from older males or for males to test strengths against age-peers (learning about their relative dominance status, instead of resources). Our results on male associations favour the testing-strengths rather than the social learning hypothesis. Similar-aged males have been shown to preferentially associate with each other and test strengths in all-male groups of other species (Villaret and Bon 1995 - Alpine ibex, Cransac *et al*. 1998 - mouflon sheep) and all-male groups in the Kabini elephant population may similarly provide a relaxed setting for old nonmusth males to test strengths with age-peers. However, young males in Kabini were sighted together as expected by chance both in female presence and absence. Similar to our results, old (30+ years) males associated more with age-peers, while the within age-class associations were as expected by chance in younger age-classes in all-male groups in the Amboseli population (Chiyo *et al*. 2011). As old males regularly enter musth, it may be more important for them than for young males to clarify their relationships when they are not in musth. It would be interesting to find out if dominance relationships formed when not in musth have a bearing on males entering into musth. Old males have also spent more time in their post-dispersal home ranges and might have formed significant associations with familiar age-peers. Male African savannah elephants in Amboseli picked familiar age-peers as sparring partners (Chiyo *et al*. 2011), and, in other species, interactions with familiar opponents tended to be less aggressive than those with unfamiliar opponents (Smith *et al*. 1999, López and Martin 2001, Wich and Sterck 2007). It would be interesting to examine the identities of sparring partners amongst males and whether familiarity plays a role in the Kabini population. Murphy *et al*. (2020) suggested that, perhaps, younger (20-30-year-old African savannah male elephants in their study) males had not yet established a consistent nonmusth home range, leading to lowered stability in association strength compared to older males. However, we found no difference between the turnover rates of young and old nonmusth males (see Supplementary Material 2); therefore, this is unlikely to be the reason for the difference observed in our six-year dataset.

We did not find any evidence for the social learning hypothesis, which suggests that the superior knowledge of experienced, older males would attract younger males to associate with them in order to learn about the location of food resources in all-male groups or about interactions with females in mixed-sex groups. Contrary to the expectations based on the social learning hypothesis, we found that young males were not sighted with old males to a greater extent than with other young males, either in female presence or absence. During the small amount of time that old and young males spent together, males were equally likely to approach the other to associate regardless of their relative ages, both in female presence and absence. Therefore, young males did not seek out old males. Although old males had higher association strengths than young males in female absence, this was due to the stronger old male-old male associations, rather than old male-young male associations.

In African savannah elephants, older males were found to be important in male networks. Older males spent more time with other males than younger males did (Poole 1982, Chiyo *et al*. 2011, Lee *et al*. 2011) and had a greater number of associates in all-male groups (Chiyo et al. 2011) in the Amboseli population. In the Samburu population, sexually inactive older males showed significant affiliation with a higher proportion of available dyads compared to sexually inactive younger males (Goldenberg et al. 2014). Old males had higher centrality in association networks based on all-male groups in Amboseli (Chiyo *et al*. 2011), although when males were classified based on their sexual state (sexually active and inactive) in the Samburu population, there was no correlation between centrality and age in sexually inactive networks and a negative correlation between centrality and age in sexually active networks (Goldenberg *et al*. 2014). In a recent study in South Africa that examined nonmusth male associations, older males showed higher stability in association strength than younger males across sampling periods; however, age of the male did not affect the stability of the identity of top ranked associates across sampling periods, eigenvector centrality, or association strength of individual males in the network (Murphy *et al*. 2020). Further, younger males preferentially associated with older males, unlike what we found in Kabini. In Amboseli, males of the 20-29 year age-class associated with older (30+) males more than expected by chance (Chiyo *et al*. 2011), and male associations seemed to facilitate social learning; males who had an older crop raider as a top associate were more likely to raid themselves (Chiyo *et al*. 2012). Older African savannah elephant males were also preferred as neighbours while associating by males of all ages, based on observations in a bull area in Okavango Delta, Botswana (Evans and Harris 2008), and have been considered analogous to the knowledgeable matriarchs of female groups in the species (McComb *et al*. 2001, Evans and Harris 2008). More generally, preferred association with, and social learning from older, more experienced individuals has been observed in the context of mating behaviour (Pereira 1988, Beecher *et al*. 1994), and foraging strategies (Rajpurohit *et al*. 1995, Biro *et al*. 2003) in other species.

The greater social role of older males in the African savannah elephant (with younger males preferentially associating with older males) compared to the Asian elephant in female absence may stem from differences in the habitats they occupy. Asian elephants occupy moister, more forested habitats, in which food is possibly more dispersed and unpredictable in space at a very local scale (but possibly more predictable at a larger spatiotemporal scale due to clear seasonality). This local heterogeneity, along with possibly limited food in a feeding patch, and greater predictability at a larger scale might make social learning as adults sub-optimal, as the habitat experienced by individual males might be different (see Boyd and Richardson 1988), and/or superfluous. We also did not find any evidence for social learning in female presence. As young male Asian elephants have been observed to obtain some mating success through opportunistic matings (see Chelliah and Sukumar 2015, Kabini Elephant Project, unpublished data), it is possible that learning in the context of reproductive behaviours occurs at a younger age. It would be interesting to examine whether there is evidence for subadult male Asian elephants socially learning from adults in mixed-sex groups. In our study population, matriarchs of female clans were not the most central individuals (Shetty 2016), which has also been suggested in Uda Walawe in Sri Lanka (de Silva *et al*. 2011). However, while female group size constraints were found to result in clans being split among small groups (Nandini *et al*. 2017), which might result in the matriarch not being central to the clan, group size constraint alone may not lead to the pattern we see amongst males. Although males also associated in small group sizes, young males did not prefer older age-class males as associates more than their age-peers, and young males did not preferentially approach old males when associations did occur. Therefore, it appears that social learning is not the primary reason for adult male associations, even accounting for limitations on group size.

### Male associations in the presence and absence of females

We found that the association network of males was non-random in female absence but random in female presence. There were no significant differences in network statistics and AI values between female presence and absence, after accounting for the different number of sightings in the two categories. Thus, the seemingly better connected network in female absence was due to the greater amount of time that males spent in female absence. Contrastingly, in African savannah elephants in Samburu, association networks of sexually inactive males were significantly denser and more clustered than those of sexually active males (Goldenberg *et al*. 2014). However, sexually active males included musth males and it is not clear to what extent this might contribute to the significant difference between the networks in Samburu.

If males were approaching female groups solely due to the presence of females and independent of the presence of other males, we had predicted that male-male associations in female presence would be random, both at an individual and at an age-class level. Our results (network degree distribution being random and all age-classes meeting each other as expected by chance) agree with these predictions. Moss and Poole (1983) observed that males in Amboseli also associated at random when in mixed-sex groups. Males in multi-male mixed-sex groups of other species have also been found to invest in affiliations with females rather than with each other (for example, mountain gorillas, see Sicotte 1994). The slightly larger male group size in female presence than in female absence could be a result of multiple males tracking the same resource (female groups).

### Restriction on all-male group size

We found a negative relationship between how many associates a male had and how much time he spent with them, indicating that all-male groups have a constraint on their group size. Males of both age-classes spent similar proportions of time in all-male groups and experienced similar all-male group sizes, which was expected if there was age-peer association among old but not young males, and a restriction on group size. Female group sizes had also been found to be constrained in the Kabini population (Nandini *et al*. 2017) and smaller compared to an African savannah elephant population (Nandini *et al*. 2018). The group size of independent sightings of all-male groups was lower in Kabini than in the Amboseli population of African savannah elephants (mean ± SD group size of all-male groups with more than one male: Kabini: 2.159 ± 0.502, Amboseli: 3.325 ± 1.995; Supplementary Material 8, see Chiyo *et al*. 2011). Low male group sizes have also been observed in other Asian elephant populations. The average group size of adult males (including solitary males) was 1.1 in Mudumalai Wildlife Sanctuary, southern India (Daniel *et al*. 1987). The maximum all-male group size was 2 in Mudumalai (Daniel *et al*. 1987), 6 in Kabini (although observed in only one sighting), and 5 in Gal Oya in Sri Lanka (McKay 1973). In contrast, the maximum group size in Amboseli was 40 (Lee *et al*. 2011). Lee *et al*. (2011) classified gregarious males (who spend most of their nonmusth time in all-male groups) as promiscuous (males who spent a lot of time with many associates) or discriminating (males who associated with fewer males for a lot of time). The presence of promiscuous males, along with much larger group sizes, suggest that there is no strong negative relationship between how many associates a male has and how strong his associations are with them in Amboseli.

There are methodological differences between the current study and the one in Amboseli. We considered 50-m as the radius for male association while Chiyo *et al*. (2011) used a 100-m radius. However, we did not find a large number of instances of males within a 100-m radius but outside a 50-m radius. Chiyo *et al*. (2011) also included males regardless of musth status/sexual state (which may artificially reduce strength of association; see Goldenberg *et al*. 2014). However, since musth males in Amboseli associated more with female groups than in all-male groups, and the reverse was true for nonmusth males (Poole 1987), it is unlikely that the removal of musth males from the data will lead to a substantial reduction in the all-male group sizes reported in Chiyo *et al*. (2011).

We expected a group size constraint to affect old males to a greater extent than young males due to greater food requirements. However, we found no effect of male age on the time spent in all-male groups. We also found no relationship between male age and the mean experienced group size, but the mean experienced group sizes were close to 1.2, precluding much further reduction in group sizes. Larger Asian elephant male groups are reported elsewhere when they raid crops in risky, but resource-rich environments (see Srinivasaiah *et al*. 2012), which would relax any constraints on group size. However, anthropogenic threat that could result in larger group sizes is a confounding factor. Intra-group feeding competition has been discussed as a constraint on group sizes in primates (see Chapman *et al*. 1995), including in all-male groups (Rajpurohit 1995, Steenbeek *et al*. 2000). Future studies on foraging in male elephants are required to find out the extent to which foraging constraints exist and affect group sizes.

### Preferred male associations and stability of associations

We did not find high correlations between associations across years, in female absence, or evidence for preferred associations among males across two week intervals in either female absence or presence. However, when we examined the males’ top associates, we found, in female absence, that the identity of the top associate was different from random for most (84%) males and a majority of the pairs (61%) had a higher AI than expected by chance, although the absolute values of these AIs were quite low (Supplementary Material 4). Thus, there were males who showed preference in their choice of top associate but did not or could not spend more time with those associates, and group size restriction may possibly play a role. Such nonrandom top associates were hardly observed in the presence of females. In all-male groups in Amboseli, most of the associations were not significant, with less than 10% of all the observed AI values being greater than that (AI=0.1) predicted under a model of random associations (Chiyo *et al*. 2011). However, similar to our findings in Kabini, older (>20 years old) adult males in Amboseli also had at least one significant top associate (Lee *et al*. 2011). Stable and significant affiliation among adult males have been observed in several other species (Packer and Pusey 1982 - lions, Connor *et al*. 2001 - bottlenose dolphins, Mitani 2009 - chimpanzees, Berghänel *et al*. 2011 - barbary macaques) but as mentioned in the Introduction, significant relationships among adult males are often thought to be a means to form coalitions to defend females. Adult male coalitions have not been observed in Asian elephants and are unlikely, given the low probability of finding a receptive female and the small sizes of female groups (Nandini *et al*. 2017). It will be interesting to explore other possible reasons for the significant top male associates we find in Kabini. It is possible that significant affiliations occur between males who are related (see Vidya and Sukumar 2005).

In summary, we show that associations among nonmusth adult male Asian elephants, although temporary, were affected by male age and the immediate presence of females. We found associations with age-peers among old males in the absence of females, likely allowing for testing strengths. There was no evidence for social learning driving male associations. There was a constraint on all-male group size, which probably contributed to the small proportion of time males spent in all-male groups. Males associated with each other at random in the presence of females. When we compared our results with those observed in African savannah elephants, we found that Kabini males spent a much smaller proportion of their time in all-male groups of smaller sizes, possibly due to the group size constraints found in Kabini. We also found no evidence for young males preferentially associating with older males in Kabini, unlike the case of African savannah elephants. We posit that the difference in the role of older males is due to the difference in the dispersion of food resources in habitats they occupy, making social learning about resources less valuable in adult male Asian elephants. Overall, while there were some similarities in male associations, ecological differences possibly result in the differences in male social structure between the two species, despite phylogenetic similarity. The relative extents to which the presence/absence of bull areas and differences in feeding competition in non-bull areas explain differences in group size and association time between adult males in the African savannah and Kabini would be interesting to examine. Differences in the nature of male associations between phylogenetically related species have been observed in other taxa: chimpanzees and bonobos show different levels of agonism (Furuichi and Ihobe 1994), macaque populations/species differ in the frequencies of male affiliations depending on group sizes and group sex ratios (Hill 1994). However, while differences between related species in overall social organisation (dolphin species: Parra *et al*. 2011, Grevy’s zebra and onager: Rubenstein *et al*. 2015) or female social organisation (colobine species: Korstjens *et al*. 2002, African savannah and Asian elephants: Nandini *et al*. 2018) that are consistent with resource distributions are known, there is little information on food resource distributions differently affecting male social organisation in related species of large mammals. Studies on the foraging ecology of male elephants are required in the future to further understand the differences in social organisation between species.

## Acknowledgments

This work was supported by the Department of Science and Technology’s (Government of India) Ramanujan Fellowship (to TNCV) under Grant No. SR/S2/RJN-25/2007 (dated 09/06/2008), Council of Scientific and Industrial Research, Government of India, under Grant No. 37(1375)/09/EMR-II and No. 37(1613)/13/EMR-II, National Geographic Society, USA, under Grant #8719-09 and #9378-13, and JNCASR. PK was supported as a Ph.D. student by the Council of Scientific and Industrial Research (No. 09/733(0152)/2011-EMR-I). This work is part of PK’s Ph.D. thesis. JNCASR also provided logistic support. The funders had no role in study design, data collection and analysis, preparation of the manuscript, or decision to publish.

We thank the offices of the PCCF, Karnataka Forest Department, and of the Conservators of Forests of Nagarahole and Bandipur National Parks and Tiger Reserves for field permits. We also thank various officials, from various PCCFs and APCCFs, to the Conservators of Forests and Range Forest Officers, to the staff of Nagarahole and Bandipur National Parks for their support across the years. We thank Krishna, Althaf, Ranga, Shankar, Gunda, Rajesh, Binu, and others for field assistance. We thank Deepika Prasad and Arjun Ghosh for help with the initial field data collection. We thank Ajay Desai for useful discussions.

