## Supplementary Material for "Effects of male age and female presence on male associations in a large, polygynous mammal in southern India"

Supplementary Material 1. Time to independence of adult male associations.

We used all the observations of adult males in the last two years of data collection (2013 and 2014) and recorded the number of changes in a male’s association within a continuous observation of that male. Only changes in the identities of the adult males in the focal male’s group and change in the focal male’s association from female presence to absence (or vice versa), were counted as changes in the focal male’s associations. Changes caused due to subadult males leaving or joining the focal male’s group were not counted in this analysis, as we were interested only in adult male-adult male associations. We summed up the number of changes in five minute intervals (all observations lasting less than five minutes were not considered for the analysis). The duration of observations used for the analyses varied from 5 to 400 minutes for musth males, and from 5 to 220 minutes for nonmusth males (mean ± SD: 37.3 ± 40.16 minutes). We examined 768 such continuous observations of males (including 24 observations of males of unknown musth status, 68 observations of musth males, and 676 observations of nonmusth males). We found that the associations of a musth males changed faster than those of nonmusth males, with an equal number of observations recording at least one change at 55 minutes of observation (11 observations with no change, and 11 observations with at least one change; Figure 1a). In the case of nonmusth males, at 80 minutes, almost half of the observations recorded at least one change (36 observations without a change, 34 observations with at least one change) and at 85 minutes, more than half of the observations had recorded at least one change (29 observations with no change, 34 observations with at least one change; Figure 1b). Therefore, we chose 80 minutes as the time at which a nonmusth male’s sighting is independent of its previous sighting.

We also examined changes in nonmusth males’ associations in each year separately to check whether they were similar across years. Based on data from 2013 (391 observations of nonmusth males), there was equal probability of an association changing or not at 75 minutes of observation (15 observations with no change, 15 observations with at least one change), indicating that 75 minutes could be considered as the time cutoff for observations to be considered independent of each other. In 2014 (285 observations of nonmusth males), there was equal probability of an association changing or not at 80 minutes of observation (21 observations with no change, 21 observations with at least one change). Thus, there was similarity across years in the time to independence of associations. We did not examine the rate of change of musth males’ associations separately in the two years as there were very few observations of musth males in 2013 (*N*=21) and the current paper deals with nonmusth males.

a) Musth males

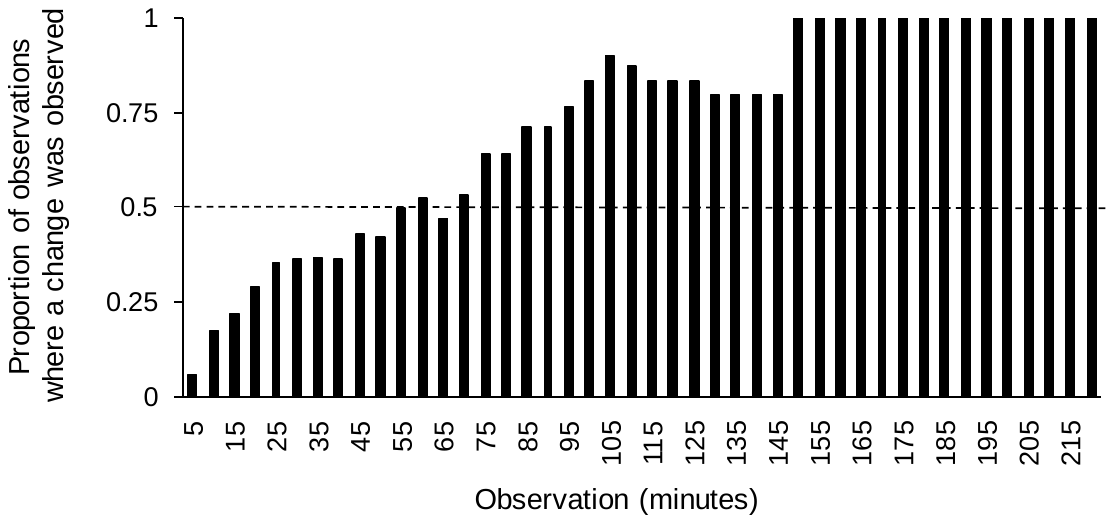

b) Nonmusth males

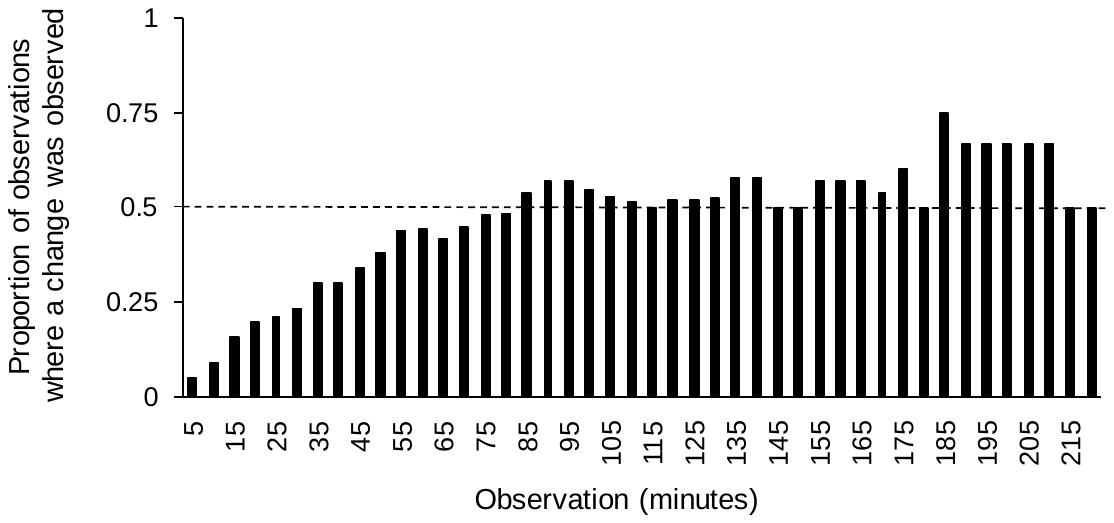

c) Nonmusth males: number of observations

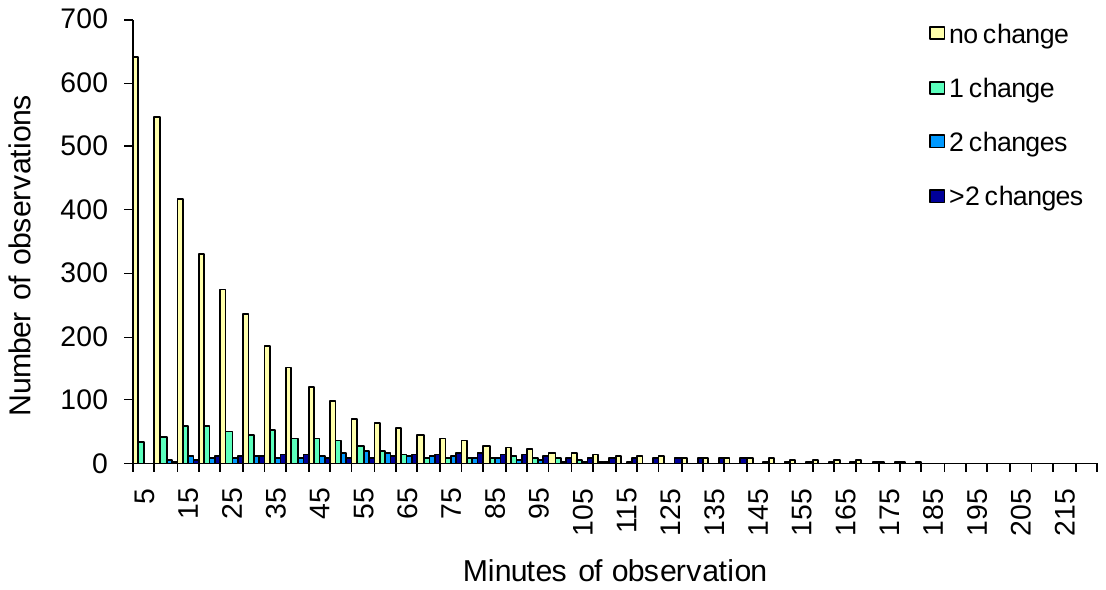

d) Nonmusth males: proportion of observations with different numbers of changes

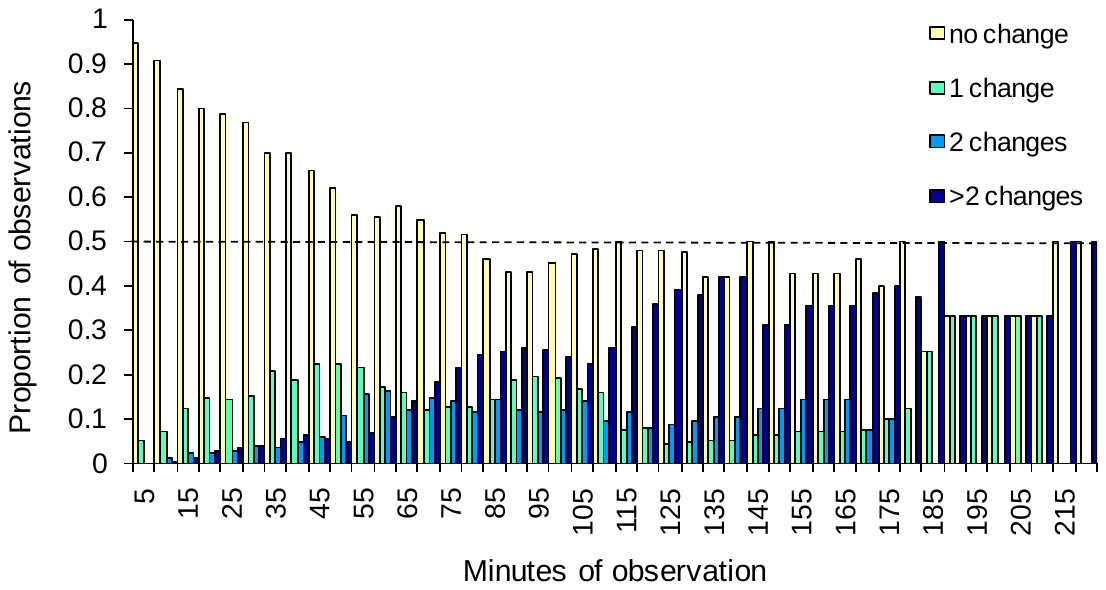

Supplementary Material 1, Figure 1. The proportion of observations that recorded at least one change in the associations of musth males (a) and nonmusth males (b), the numbers of observations with different numbers of changes in the associations of nonmusth males (c), and the proportions of observations with different numbers of changes (d), at different durations of observation, using data from 2013 and 2014.

As mentioned in the main text, using an 80-minute cutoff, we found 14 nonmusth males who were sighted ten times or more in female presence (mean ± SD of number of sightings: 21.4 ± 13.40 sightings, maximum: 65 sightings, minimum: 13 sightings), and 32 nonmusth males who were sighted ten times or more in female absence (mean ± SD of number of sightings: 66.2 ± 54.85 sightings, maximum: 216 sightings, minimum: 10 sightings). There were only 12 common males who were sighted ten times or more both in female presence (mean ± SD of number of sightings: 22.7 ± 14.10 sightings, maximum: 65 sightings, minimum: 13 sightings), and in female absence (mean ± SD of number of sightings: 108.7 ± 50.79 sightings, maximum: 216 sightings, minimum: 46 sightings).

Supplementary Material 2. Number of males identified.

We identified 96 adult males in all (based on all sightings in which all adult males were aged and identified). Out of these, 83 males were sighted when they were not in musth. Only 44 nonmusth males were seen in the presence of females and 81 nonmusth males were seen in the absence of females. Since this is an open population, it is natural for new elephants to arrive into the area due to dispersal. The cumulative number of identified adult males is shown in Figure 1 below.

We also examined the turnover rates for old and young nonmusth males sighted across the years (2011-2014, Figure 1b), and found that the age-class composition of males sighted in the first half of this period (2011-2012) was not different from that of males sighted for the first time in the second half of the study period (2013-2014; 2 x 2 *G*-test of independence: *G*_adj_=0.770, *df*=1, *P*=0.380). This suggests that the turnover rates of old and young males were not different across this study period.

| a)  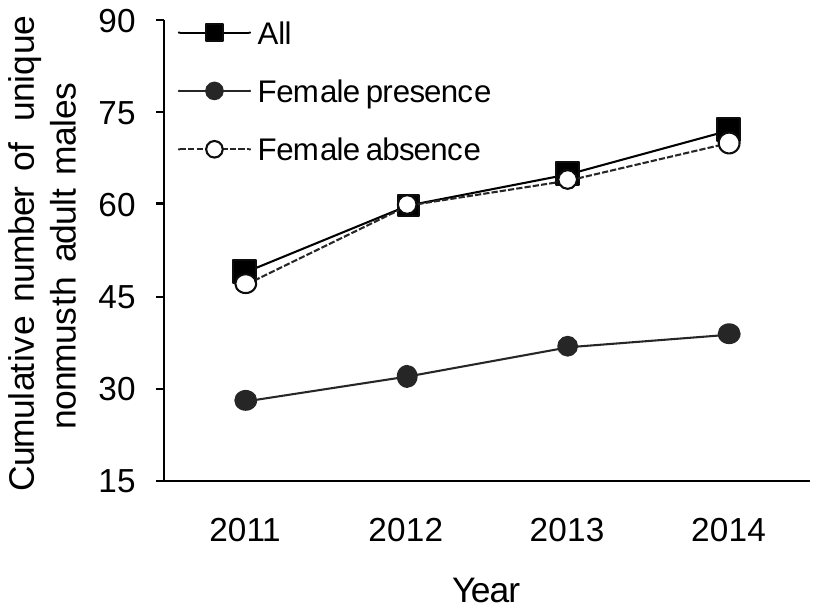 | b)  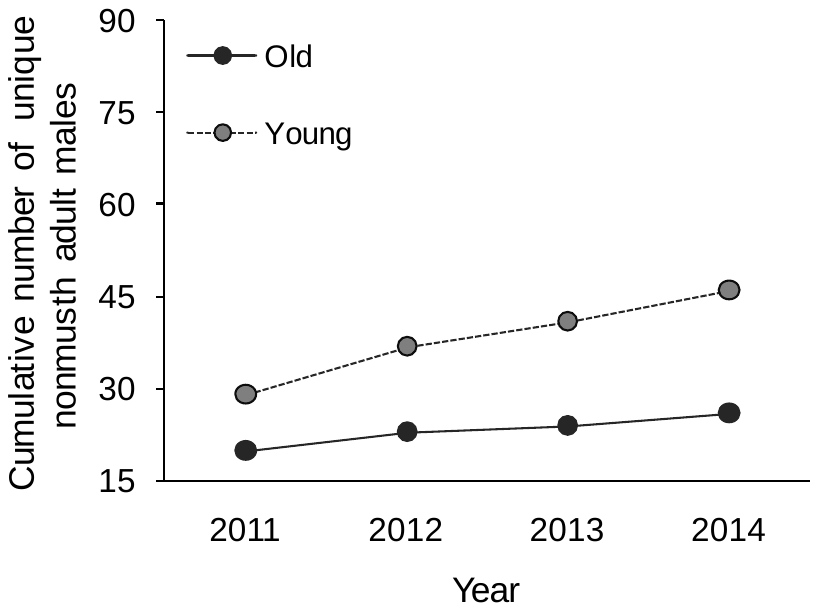 |
| --- | --- |

Supplementary material 2, Figure 1. Cumulative numbers of a) identified nonmusth adult males overall, in female presence, and female absence, and b) nonmusth old and young adult males sighted across years.

Supplementary material 3. Males sighted with other males of the same and different age-classes in female presence and absence, using all identified males.

As mentioned in the Methods and Results, we compared the observed male associations in female presence and absence with those obtained by permuting males within the female presence or female absence datasets. While results based on sightings of males sighted ten times or more are shown in the Results, we also carried out this analysis on the dataset of all identified adult males. The results from this analysis are below. Randomising males between sightings in female presence, we found that males of all age-classes were sighted together as expected by chance (old-old: observed: 5 sightings, permuted (2.5 and 97.5 percentiles): 3.8 (1, 7) sightings, *P*=0.169; old-young: observed: 20 sightings, permuted: 19.1 (13, 25) sightings, *P*=0.316; young-young: observed: 23 sightings, permuted: 23.0 (17, 28) sightings, *P*=0.436; Figure 1a below). In female absence, old males were sighted together more than expected by chance (observed: 50 sightings, permuted: 28.0 (20, 37) sightings, *P*<0.001), old and young males were sighted together less than expected (observed: 59 sightings, permuted: 75.8 (65, 87) sightings, *P*=0.003), and young males were sighted together as expected by chance (observed: 47 sightings, permuted: 49.3 (39, 59) sightings, *P*=0.317; Figure 1b below). Thus, the results obtained based on all identified males were similar to the results shown in the main text based on only the common males.

| a) Female presence  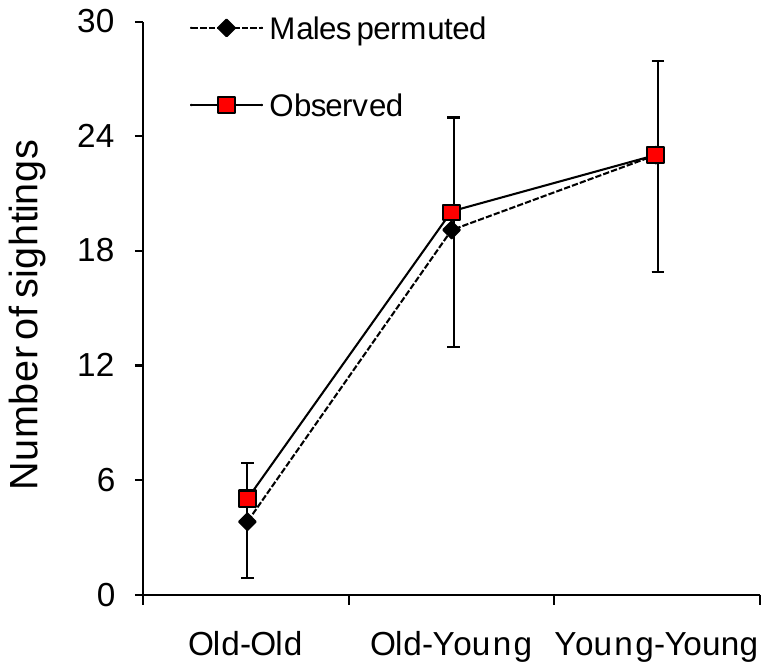 | b) Female absence  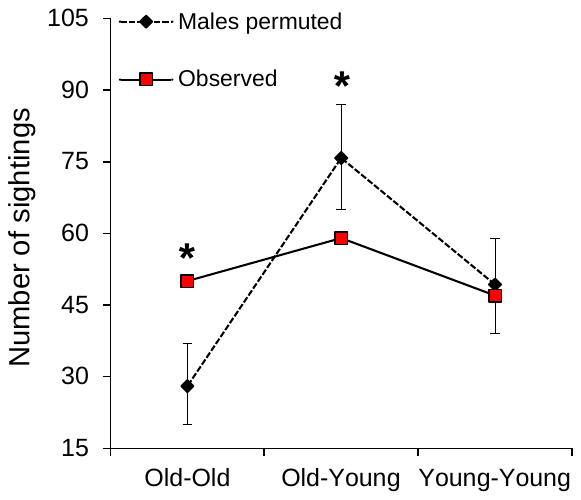 |
| --- | --- |

Supplementary material 3, Figure 1. Permuted and observed numbers of times adult males of the same and different age-classes were sighted together in a) female presence and b) female absence. Comparisons where the observed value was significantly different from those of the permuted datasets are marked with an asterix. Old males are >=30 years old and young males are 15-30 years old. Please note that the Y axis scales are different in the two panels.

Supplementary Material 4. Distribution of AI values in female presence and female absence.

The distributions of AI values between identified males seen in female presence and female absence are shown below (Figure 1). The distribution of AI values between males and their top associate in female absence which were significantly higher than expected (*N*=19, mean (SD): 0.032 (0.015)) are shown in Figure 1c. There were hardly any significant top associates in female presence.

| a)  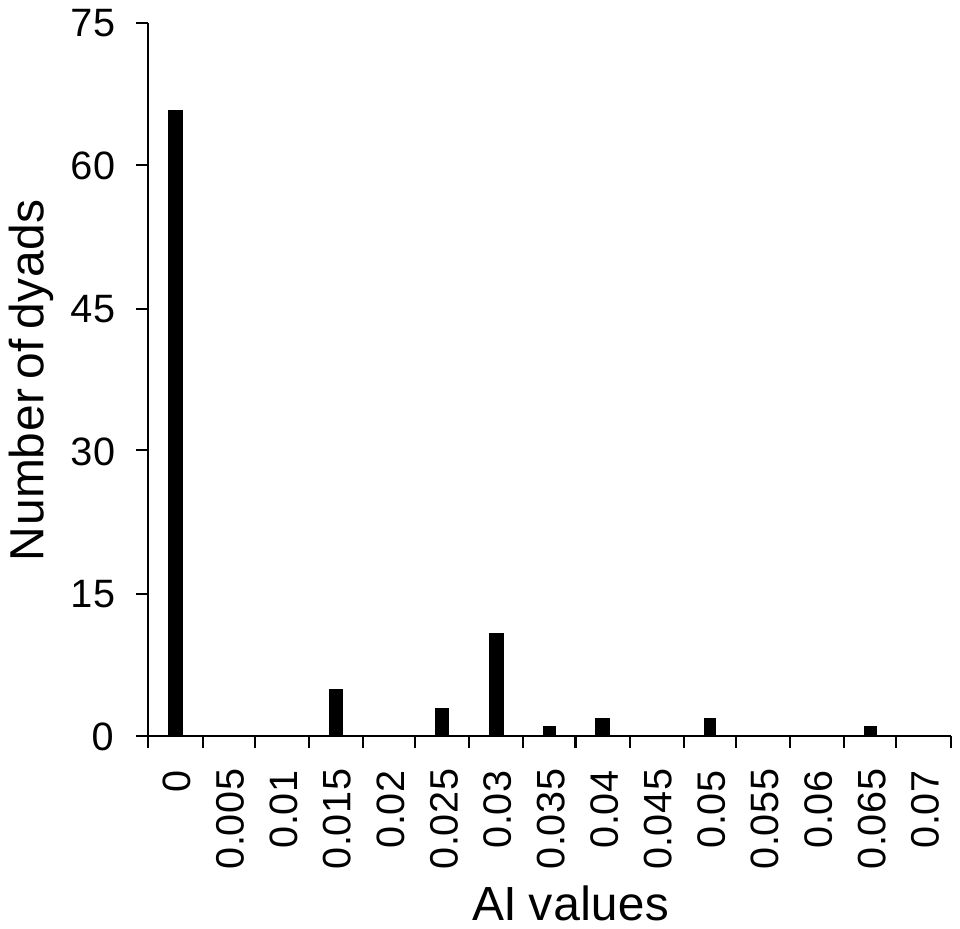 | b)  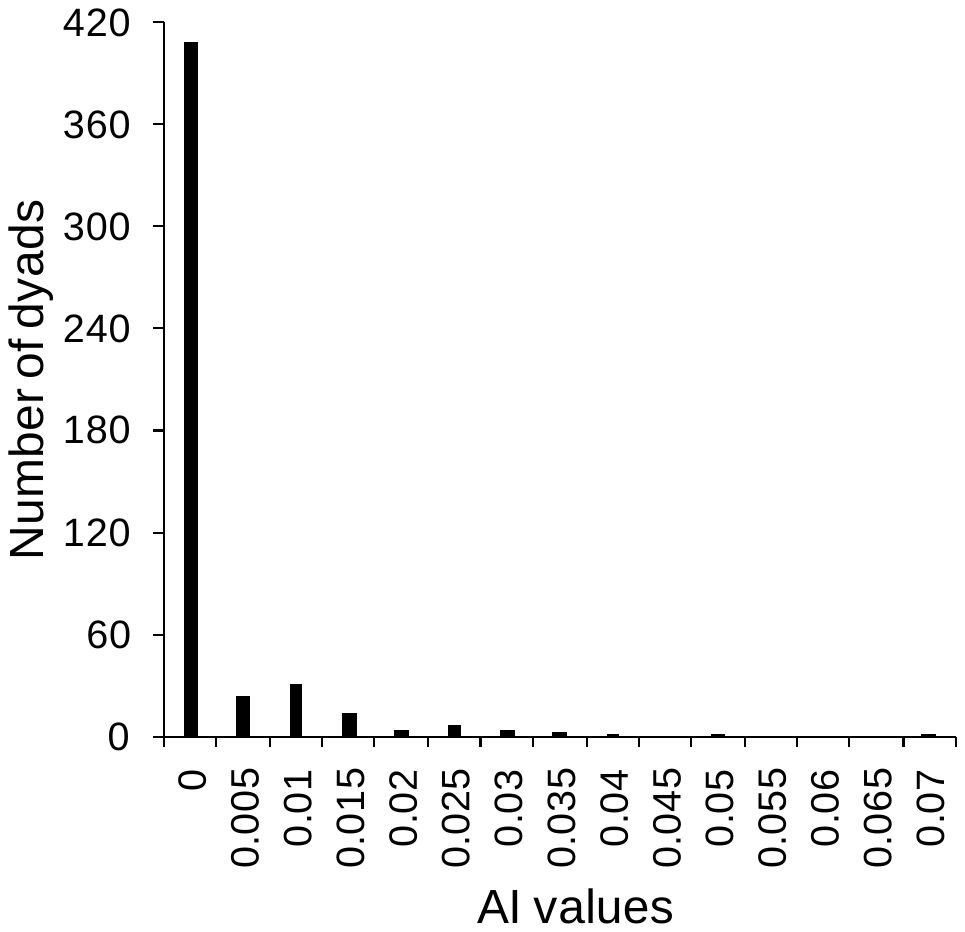 |
| --- | --- |
| c)  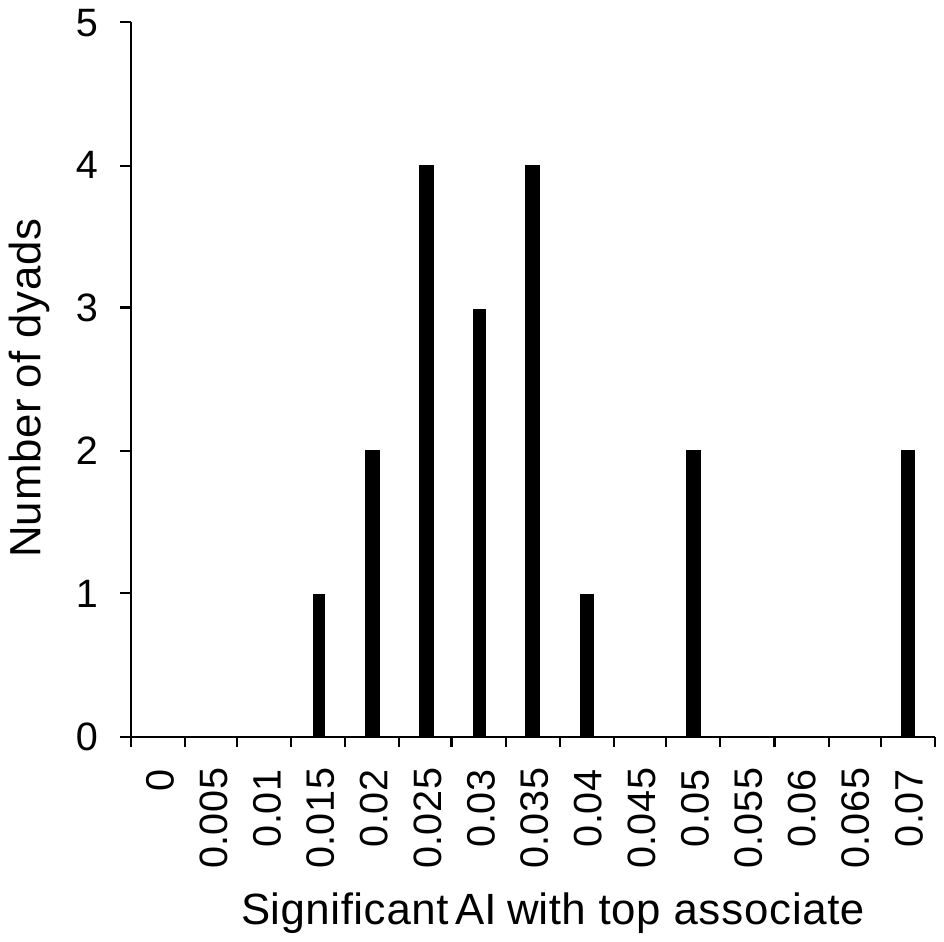 |  |

Supplementary Material 4, Figure 1. AI values among males sighted 10 times or more in a) female presence (*N*=14) and b) female absence (*N*=32), and AI values between 19 males and their significant top associates in female absence (c). Please note that the Y axis scales are different across panels.

Supplementary Material 5. Degree distributions of common males in female presence and female absence.

We compared the degree distributions of observed male association networks with Poisson distributions, expected from Erdös-Rényi random networks (Erdös and Rényi 1960). The observed degree distribution in female presence (based on 14 males sighted ten times or more) was not significantly different from Poisson expectation (*χ^2^*=6.122, *df*=8, *P*=0.633), while the observed degree distribution in female absence (based on 32 males sighted ten times or more) was significantly different (*χ^2^*=184.647, *df*=15, *P*<0.001; Figure 1 below).

| a)  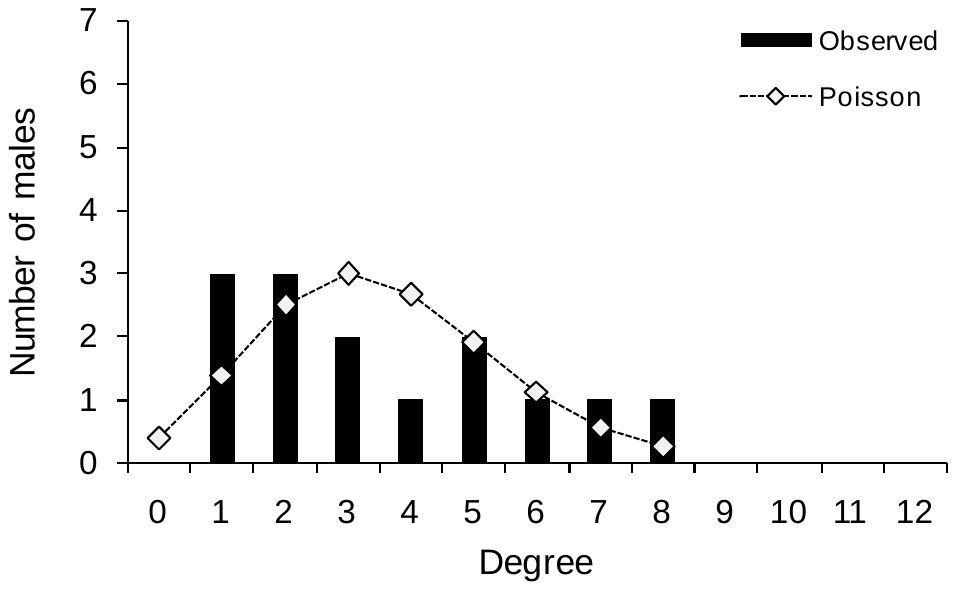 | b)  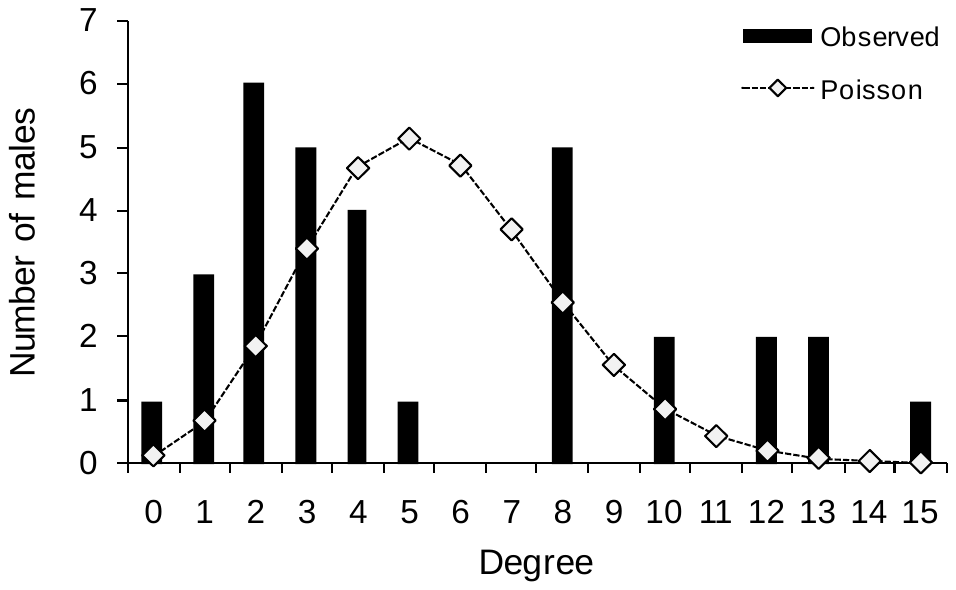 |
| --- | --- |

Supplementary Material 5, Figure 1. Observed (bars) and expected (lines) degree distributions of male association networks in a) female presence and b) female absence.

Supplementary Material 6. Permutation tests for preferred associations.

We used SOCPROG 2.6 to perform permutations tests to check for preferred associations across 14-day sampling intervals (*permute associations within samples*). This method accounts for differences in gregariousness. We used 10,000 permutations with 10,000 flips per permutation for this test and examined the observed and random SD and CV of AI (mean AI is not meaningful in this test). If the observed SD and CV of AI were significantly greater than 95% of the corresponding values in the permuted datasets, then there is long-term preferential associations among the males across the sampling intervals. The results of the permutation tests are tabulated below (Table 1). There were no statistically significant values (all *P*>0.05), indicating that adult males did not show preferred associations.

Supplementary Material 6, Table 1. Observed and random values of statistics and *P* values from the permutation test for preferred associations in female presence and in female absence, using 10,000 permutations and 10,000 flips per permutation. The number of identified males in each category is shown.

| Category | Statistic | Observed value | Ave. random value using 10000 flips | *P* (1-sided)  (10000 flips) |
| --- | --- | --- | --- | --- |
| Adult males | Mean AI | 0.0031 | 0.0030 | - |
| in female | SD of AI | 0.0166 | 0.0155 | 0.1747 |
| presence; *N*=44 | CV of AI | 5.2550 | 5.1318 | 0.3289 |
|  | Mean non-zero AI | 0.0624 | 0.0596 | 0.1784 |
|  | SD of non-zero AI | 0.0421 | 0.0371 | 0.2476 |
|  | CV of non-zero AI | 0.6752 | 0.6204 | 0.3582 |
| Adult males | Mean AI | 0.0010 | 0.0010 | - |
| in female | SD of AI | 0.0071 | 0.0071 | 0.3524 |
| absence; *N*=81 | CV of AI | 7.2425 | 7.0756 | 0.1246 |
|  | Mean non-zero AI | 0.0313 | 0.0309 | 0.3330 |
|  | SD of non-zero AI | 0.0260 | 0.0250 | 0.2385 |
|  | CV of non-zero AI | 0.8282 | 0.8084 | 0.2362 |

Supplementary material 7. Mantel test results of correlations between association index matrices of consecutive years.

We carried out Mantel tests of matrix correlations between AI matrices of consecutive years in female absence, using common males (nonmusth males sighted five times or more in each of the two years being compared). Only one of the three comparisons yielded a significant correlation, which was low (Table 1 below). Similar comparisons could not be made between males sighted in female presence as there were very few common nonmusth males across years in this category.

Supplementary material 9, Table 1. Mantel test results (based on 5000 permutations) from comparing AI matrices of common males in consecutive years in female absence.

| Comparison |  | Female absence | | |
| --- | --- | --- | --- | --- |
|  | *N* | *R* | *R^2^* | *P* |
| 2011-2012 | 18 | -0.030 | 0.001 | 0.666 |
| 2012-2013 | 18 | -0.066 | 0.004 | 0.817 |
| 2013-2014 | **16** | **0.257** | **0.066** | **0.014** |

Supplementary Material 8. Adult male group sizes in all-male groups in Kabini and Amboseli.

We examined the adult male group sizes in all-male groups in Kabini and Amboseli. Over decades of observation, all-male group sizes in Amboseli were found to vary form 2-40 (Lee *et al*. 2011). Chiyo *et al*. (2011), based on three years of data from Amboseli, found the mean group size (number of adult males) in all-male groups to be (mean ± SD) 3.325 ± 1.995 (*N*=939 groups). We compared the mean all-male group sizes (using independent sightings) in Kabini (mean ± SD: 2.159 ± 0.502, *N*=138) with those in Amboseli using Welch’s two-sample test (Welch 1937, see Fagerland and Sandvik 2009), and found adult male groups in Amboseli to be significantly larger than those in Kabini (Welch’s two-sample test: *U*=14.969, *fu*=846.050, *P*<0.001). Thus, males in the Kabini population not only spent much less time in all-male groups than in the Amboseli African savannah elephant population, but, also formed groups of smaller sizes when they did associate in all-male groups.
